# Mechanistic Insights into Keratin Degradation by *Onygena corvina*

**DOI:** 10.1101/2025.07.09.663871

**Authors:** Siddhi Pavale, Clémentine Isembart, Volha Shapaval, Tina R Tuveng, Sabina Leanti La Rosa, Vincent G.H. Eijsink

## Abstract

Keratin-rich byproducts from the poultry, textile, and leather industries pose a significant challenge for sustainable waste management due to their highly recalcitrant nature. While microbial degradation may offer a viable solution, the mechanisms underlying keratin breakdown remain largely unexplored. In this study, we employed a high-resolution proteogenomic approach to characterize the keratinolytic machinery of *Onygena corvina*, a non-pathogenic saprophytic fungus. Using a membrane agar plate method with insoluble substrates, we obtained secretomes enriched in secreted and substrate-bound proteins during growth on α- and β-keratin-rich substrates, specifically wool and feather meal. Our findings reveal that *O. corvina* has a richer proteolytic machinery than previously reported, including enzymes that are used across keratin types, as well as enzymes that are specifically targeted to either α- or β-keratin. In addition to proteases, the secretomes contain numerous other proteins, including cell wall-modifying enzymes, oxidoreductases, esterases, phosphatases, and sialidases that are involved in the deconstruction of keratin. We propose that these additional enzymes destabilize keratin through a combination of mechanical keratinolysis, removal of post-translational modifications, reduction of disulfide bonds, and cleavage of isopeptide bonds, thereby enhancing proteolytic accessibility. Interestingly, keratin degradation by *O. corvina* was most efficient when using mixed substrates containing both feather and wool meal. These novel insights into the keratinolytic system of *O. corvina* underscore the importance of considering synergistic enzyme interactions when developing biotechnological approaches for valorization of keratin-rich by-products.

## Introduction

Keratin is a fibrous structural protein that is predominantly found in animal-derived hard tissues such as feathers, wool, hair, horns, and hooves (1). Annually, over 30 billion tons of keratin-rich byproducts are generated worldwide, primarily from the poultry, textile, and leather industries (2, 3). Among these, feathers and wool constitute the major sources of keratin-rich biomass, offering significant potential for the conversion into biodegradable materials, biofertilizers, animal feed, and bioactive peptides (4). Despite this, efficient valorization of keratinous byproducts remains a challenge due to the highly recalcitrant structure of keratin, which limits its degradation by conventional proteases. Keratin is composed of intermediate filaments that are bundled into fibrous structures, and its structure is stabilized by disulfide bonds, hydrogen bonding and hydrophobic interactions. α-keratin, which is rich in α-helices and the predominant form in wool and hair, exhibits greater resistance to enzymatic degradation than β-keratin, which is rich in β-strands and the primary constituent of feathers (5–7).

Conventional keratin-rich byproduct processing strategies include thermal, chemical, and mechanical treatments, which are energy-intensive and environmentally detrimental, while yielding products of limited value (4). As a sustainable and economically viable alternative, microbial degradation of keratin has emerged as a promising approach. Various microorganisms, including bacteria, fungi, and actinomycetes, have demonstrated keratinolytic capabilities (8). The first report of microbial keratin degradation dates back to the late 19^th^ century with the discovery of keratinolytic activity in a saprophytic fungus, *Onygena equina* (9, 10). Since then, extensive studies have sought to identify and characterize keratin-degrading enzymes, particularly proteases. It has been suggested that degradation of keratin requires a minimum of three distinct types of proteases, including *endo*-acting, *exo*-acting, and oligopeptide-acting enzymes (11–13). However, proteases alone are insufficient to efficiently degrade native keratin due to its extensively cross-linked structure, which limits enzyme accessibility. Indeed, effective keratin degradation requires the cooperative action of multiple enzymes, including disulfide bond-reducing enzymes, such as disulfide reductases, that loosen keratin’s cross-linked structure (14–16). Studies of dermatophytes such as *Trichophyton rubrum* have shown that these fungi combine invasive growth through specialized hyphal structures with a multi-enzyme strategy involving a diverse repertoire of secreted proteases, oxidoreductases, and other hydrolases, to infect keratinized structures like the epidermal stratum corneum, hair and nails (17, 18).

Despite these insights, the development of cell-free enzyme preparations for keratin valorisation has largely focused on proteases, overlooking the broader enzyme repertoire that may be needed to achieve efficient depolymerization (19). Furthermore, while dermatophytes have been extensively studied, the mechanisms underlying keratin degradation in non-pathogenic fungi remain largely unexplored. One such fungus is *Onygena corvina*, a member of the Onygenales order, known for its ability to grow directly on keratinous substrates (20). Unlike dermatophytes, which invade host keratin, *O. corvina*is a saprophyte that thrives on decaying keratinous materials, suggesting it has evolved an efficient extracellular enzymatic system for keratin utilization (21).

In this study, we have employed a proteogenomic approach to gain comprehensive insight into the keratinolytic machinery of *O. corvina*. Our findings provide evidence of an intricate keratin degradation system that extends well beyond proteolysis and differs between wool and feathers. The identification of various cell wall-modifying enzymes, oxidoreductases, esterases and phosphatases suggests hyphal penetration of the substrate and an important role for non-proteolytic enzymes that weaken the keratin structure. These results shed light on the complexity of microbial keratin degradation and show the need for a holistic approach to bioprocessing of keratin-rich biomass.

## Results and Discussion

### Proteolytic potential of *O. corvina* based on genome and secretome data

For comprehensive proteomics studies, a high-quality reference database is essential (22). Since the publicly available genome for *O. corvina* was highly fragmented (GCA_000812245.1; 521 contigs), we used a combination of short-reads (Illumina) and long-reads (Oxford Nanopore Technology) sequencing, to obtain a highly contiguous assembly (23). The new genome assembly contains 13 contigs, yielding a genome size of 21.8 Mb and a completeness of 98.97%. Annotation of protein-coding genes was performed through an integrated approach that combined de novo predictions, predictions based on homology, and transcriptome-based (RNA-Seq) evidence. This workflow yielded 7232 protein sequences, which were used to construct a custom database for subsequent proteomics studies.

Among these 7232 proteins, 284 (3.9%) were predicted to be secreted based on the presence of a signal peptide and absence of transmembrane domains. A “Homology to Peptide Pattern” (Hotpep) search (24) identified 158 genes encoding proteases, whereas only 73 were predicted in the previous genome assembly (20). Of these 158 putative proteases, 59 (37.3%) were predicted to be secreted and distributed across five major MEROPS (25) families: serine (32; S), metallo (20; M), cysteine (1; C), threonine (2; T), and aspartic (4; A). Only a subset of these secreted proteases, 30 in total, primarily belonging to the serine and metalloprotease families (Figure 1), were expressed by *O. corvina* during growth on casein, feather or wool meal, as discussed in detail in the sections below.

**Figure 1:**
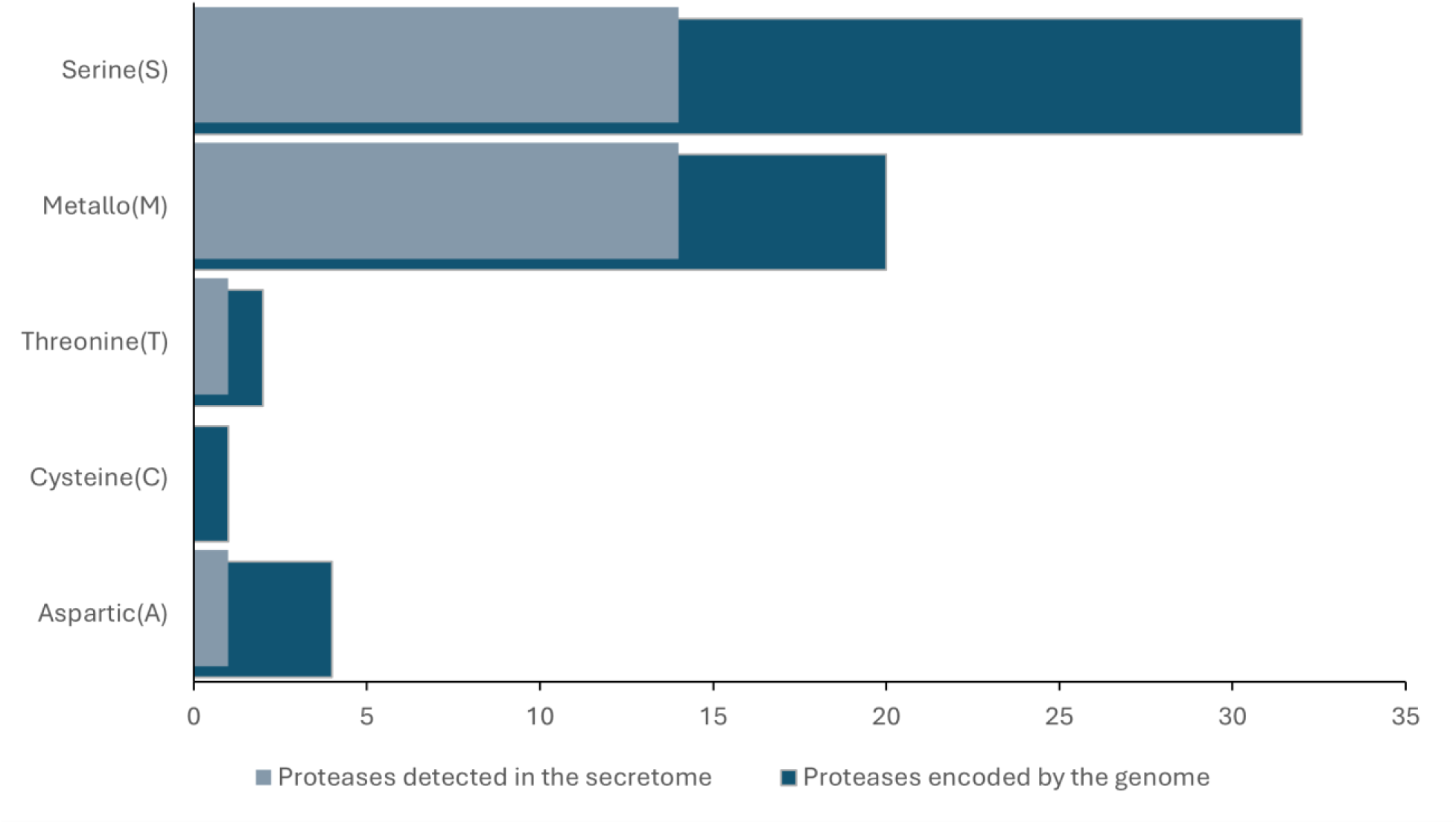
Putatively secreted proteases encoded by the *O. corvina* genome and those detected in the secretome. The bar chart displays the distribution of proteases, predicted to be secreted, across five major MEROPS families: serine (S), metallo (M), threonine (T), cysteine (C), and aspartic (A). Dark blue bars represent all proteases predicted to contain a signal peptide in the *O. corvina* proteome, while light blue bars indicate the subset of proteases detected in the secretome of *O. corvina* cultured on keratin-rich substrates (feather and wool meal) and casein.

### Secretome composition across the different substrates

To assess the ability of *O. corvina* to degrade various protein-rich sources, the fungus was grown on agar plates consisting of a minimal medium (see methods) supplemented with feather meal, wool meal or casein as the sole source of carbon and nitrogen, where casein represents an easily degradable protein source. Although *O. corvina* displayed similar radial growth on all three substrates, a clearance zone around the colonies was detected within 2 days of inoculation on casein, and after 8 days on feather and wool meal (Figure S1). This delayed detection of clearance zones during growth on recalcitrant keratin-rich substrates aligns with previous reports (26, 27).

To identify secreted proteins, samples were collected from the bottom agar of membrane plates containing one of the three substrates at 1, 2, and 3 days after inoculation. Of note, samples collected after day 3 were excluded from analysis, due to the low number of detected proteins. High-resolution LC-MS/MS analysis, followed by protein quantification using the topN algorithm (28), showed high reproducibility, with most Pearson correlation coefficients being well above 0.8 and, in more than half of the cases, above 0.9 (Figure S2-S4). Proteins detected in at least two of the three replicates for at least one of the three substrates, 154 in total (Table S1), were considered for further analysis. Among these, 80 proteins had a predicted signal peptide, of which 7 also contained transmembrane domains, yielding 73 putatively secreted proteins (Table S2 and Figure S5). An average of 52.2% proteins identified across the secretomes from casein and keratin-rich substrates, were predicted to be secreted (Table S3). Given that only 3.9% of the total *O. corvina* proteome is putatively secreted, the membrane plate method effectively captured and significantly enriched for extracellular proteins.

Focusing on the 73 putatively secreted proteins, the composition of the secretomes differed markedly between keratin-rich substrates and casein. Only 13 proteins were detected in the casein secretome, all of which were also present in the feather and wool secretomes, which contained 59 and 67 proteins, respectively (Figure S5). The higher number of proteins secreted on keratin-rich substrates likely reflects the greater structural complexity of keratin, necessitating a more extensive enzymatic repertoire for degradation. Of the 60 proteins found exclusively in the keratin-grown secretomes, 44 were shared between the wool and feather meal secretomes (Figure 2A), suggesting a common enzymatic machinery for the breakdown of both α- and β-keratin, alongside adaptations specific to each substrate.

**Figure 2:**
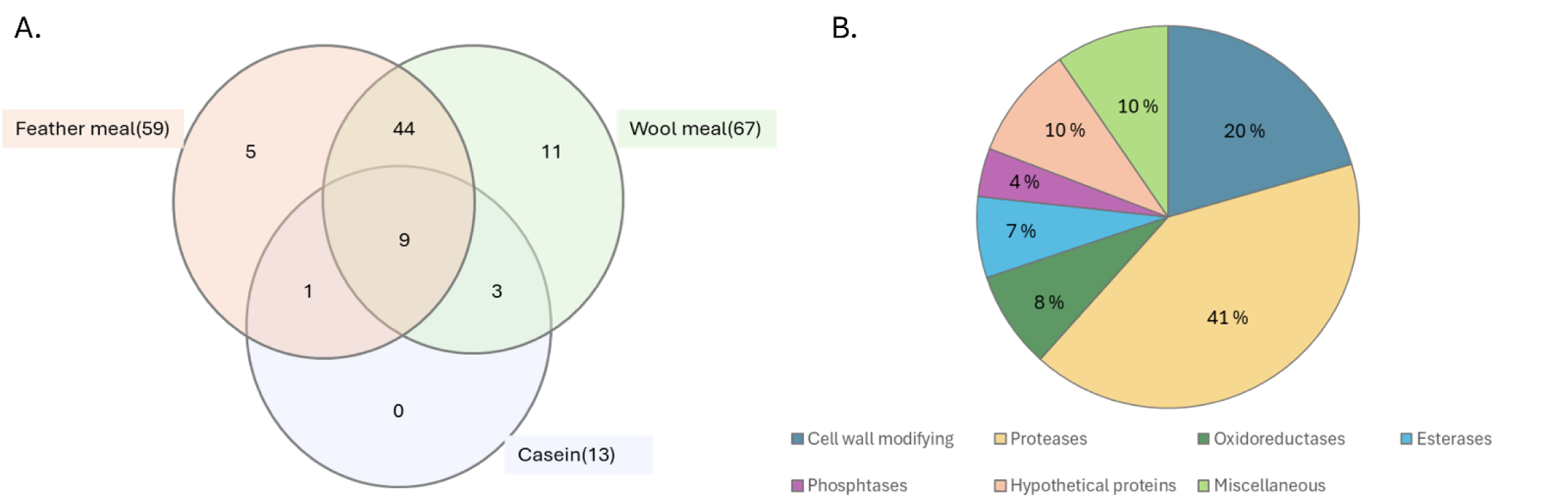
Number and functional annotation of 73 putatively secreted proteins detected in *O.corvina* secretomes. (A) Venn diagram representing the distribution of 73 putatively secreted proteins in the secretome during growth on feather meal, wool meal, or casein. The total number of detected proteins for each substrate in indicated in brackets. (B) Functional annotation of the 73 putatively secreted proteins. The figure depicts the proportion of different classes of proteins detected in the *O. corvina* secretome, irrespective of the substrates used for growth. The proteins were annotated using InterPro Scan, UniProt and BLAST analysis, followed by manual curation into the functional categories. Information regarding the nature and detection levels of these proteins can be found in Table S2 and Figure S5.

Functional annotation revealed that proteases represent the largest category, making up 41% of the secretome (30 of 73 putatively secreted proteins), while other groups included cell wall-modifying enzymes, oxidoreductases, esterases, phosphatases, as well as hypothetical proteins and proteins with miscellaneous functions (Figure 2B). Notably, hypothetical proteins with unknown functions accounted for 10% of the detected, putatively secreted proteins. The casein secretome was dominated by eight proteases alongside three hypothetical proteins, one esterase, and one oxidoreductase (Figure S5). Overall, the presence of 60 putatively secreted proteins unique to the keratin-grown secretomes highlights their likely roles in keratin degradation.

### Secretion of proteases

Growth of *O. corvina* on keratin-rich substrates led to the secretion of a diverse range of proteases, primarily from the serine (S) and metalloprotease (M) families. Out of the 158 putative proteases encoded by the *O. corvina* genome, 35 were detected by proteomic analysis following growth on feather meal, wool meal, and/or casein. Of these, 32 proteases contained a predicted signal peptide, including 2 with accompanying transmembrane domains, yielding 30 putatively secreted proteases. The remaining 3 proteases, lacking a predicted signal peptide, were classified into the M1, M3 and M49 metalloprotease families.

A heat map of the 32 proteases with a predicted signal peptide, revealed five distinct clusters (Figure 3). Cluster I comprises high-abundance proteins detected with all three substrates. Of note, several of these proteases were more abundant and found at more time points for wool and feather meal, compared to casein. Cluster II contains high-abundance proteases detected exclusively during growth on the keratin-rich substrates. Clusters III, IV, and V comprise a wide variety of proteases with relatively low abundance. The proteases in Cluster IV were detected with both keratin-rich substrates, whereas Clusters III and V contain proteases unique to wool and feather meal, respectively.

**Figure 3:**
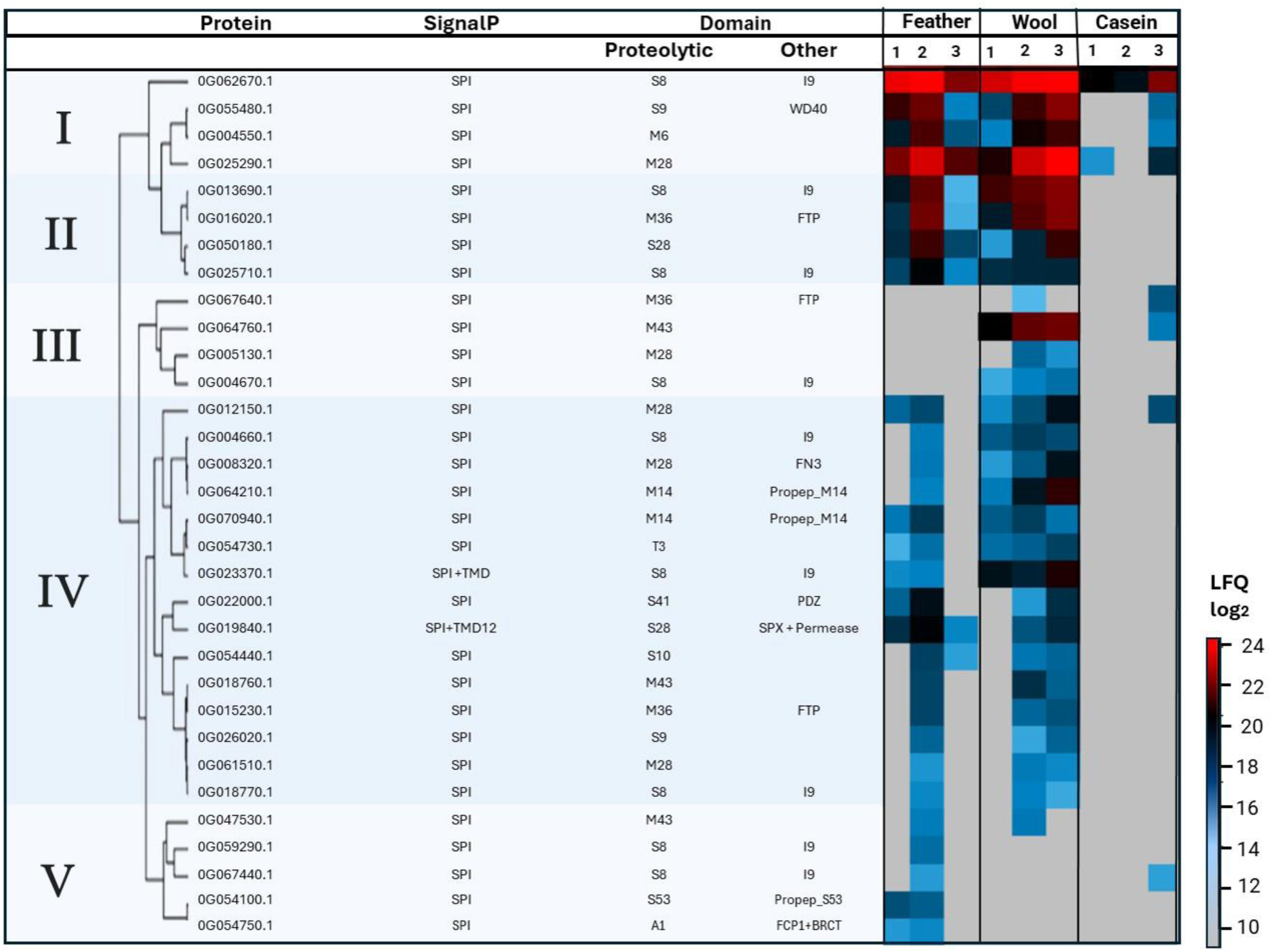
Heat-map representation for 32 detected proteases with a predicted signal peptide (SPI). The figure shows Protein ID, SignalP 5.0 and TMHMM2.0 predictions (SPI – standard secretory peptide, TMD– Transmembrane domain), the protease class, the presence of additional domains and the expression pattern (as log_2_ LFQ) of the proteases. Every row represents one protein, and the colors in the heatmap depict protein abundance (average of three replicates) during growth on feather meal, wool meal or casein at day 1, 2 or 3. The scale of the heatmap ranges from high abundance (red color) to low abundance (light blue color). Grey color indicates that the protein was not detected. Proteases are divided into five hierarchical clusters based on protein abundance and expression pattern (Cluster I-V). The domains, belonging to peptidase and other families, were assigned using Interpro Scan. Abbreviations: I9, Peptidase Inhibitor I9; WD40, 40 amino acid motif with terminal Trp-Asp (W-D) dipeptide; FTP, Fungalysin/Thermolysin Propeptide; FN3, Fibronection III domain; Propep_M14, M14 propeptide; PDZ, domain originally identified in PSD95, Dlg, and ZO-1 proteins; SPX, domain originally identified in SYG1, Pho81, and XPR1 proteins; Propep_S53, S53 propeptide; FCP1+BRCT, phosphatase and BRCA1 C-terminus domains.

Cluster I includes four highly abundant proteases that were detected with all substrates and, in the case of the keratin-rich substrates, across all time points. Two of these belong to the S8 and M6 families of *endo*-proteases, while the others are classified within the S9 and M28 families of *exo*-proteases. The presence of both *endo*- and *exo*-acting proteases suggests a synergistic mechanism for keratin degradation. Such synergism has been observed in an earlier study of *O. corvina* growing on pig bristles, in which a synergy between G062670.1 (corresponding to GenBank Accession: KP290860.1; S8 protease) and G025290.1 (corresponding to GenBank Accession: KP290838.1; M28 protease) was reported (20). The two *exo*-acting proteases in Cluster I belong to the S9 (G055480.1) and M28 (G025290.1) families, which are different and exhibit distinct substrate preferences. The S9 family includes proteases which cleave substrates from the N-terminus, typically preferring those with a proline in the penultimate position (25, 29). On the other hand M28 aminopeptidases remove N-terminal residues with a preference for large hydrophobic amino acids like leucine (25). Interestingly, while M28 peptidases cannot accommodate proline in the penultimate position (30), S9 peptidases preferentially cleave when such a proline is present, which indicates that *O. corvina* secretes a complementary set of *exo*-proteases to maximize degradation efficiency. Since Cluster I proteases were also detected on casein, they may not be specifically induced by keratin but rather be proteases with general functions that are triggered by the mere presence of protein and peptides derived thereof. This aligns with a previous study showing that G062670.1 (corresponsing to GenBank Accession: KP290860), the most abundant protease detected in this study, is an enzyme with high cleavage efficiency and no apparent cleavage specificity (31).

Cluster II is enriched in *endo*-proteases from the S8 and M36 families, along with a carboxypeptidase from the S28 family, all of which were detected exclusively during growth on keratin-rich substrates. The involvement of M36 and S8 proteases in degradation of native keratin has been well documented (11, 12, 32). The S28 family consists of carboxypeptidases that hydrolyze prolyl bonds, making these enzymes particularly relevant for the degradation of keratin, which is rich in proline (12, 33). The exclusive detection of Cluster II proteases on keratin-rich substrates suggests that they are keratin-specific; by targeting otherwise recalcitrant parts of the keratin, they may complement the broad-specificity proteases in Cluster I.

Cluster IV, the cluster with the highest number of proteins, contains less abundant proteases detected exclusively during growth on keratin-rich substrates. It includes members from a diverse range of protease families, including the serine protease (S8, S9, S14), metalloprotease (M14, M28, M36, M43), and threonine protease (T3) families. A unique feature within this cluster is the presence of a C-terminal FN3 (fibronectin III) domain in G008320.1, a M28 family protease (Figure 4). FN3 domains share topological similarity with immunoglobulin domains and exhibit diverse ligand-binding properties, potentially contributing to substrate specificity (34, 35). Another protease in this cluster, G022000.1, belonging to the S41 family, contains a PDZ domain (Figure 4), which is a peptide-binding domain that may facilitate substrate binding (36, 37). Studies of a *Vpr* serine protease from *Bacillus cereus* DCUW showed that deletion of its C-terminal PDZ domain led to loss of feather degradation ability, while retaining activity against simpler substrates like casein and gelatin (38). These observations suggest that Cluster IV proteases may exhibit activity toward specific peptide sequences within keratin, acting in concert with Cluster II proteases to disrupt native keratin structures. Of note, based on previous studies on *Trichophyton rubrum* (39), the M14 carboxypeptidases (G064210.1,G070940.1) in Cluster IV likely require activation by subtilisin-like proteases from the S8 family.

**Figure 4:**
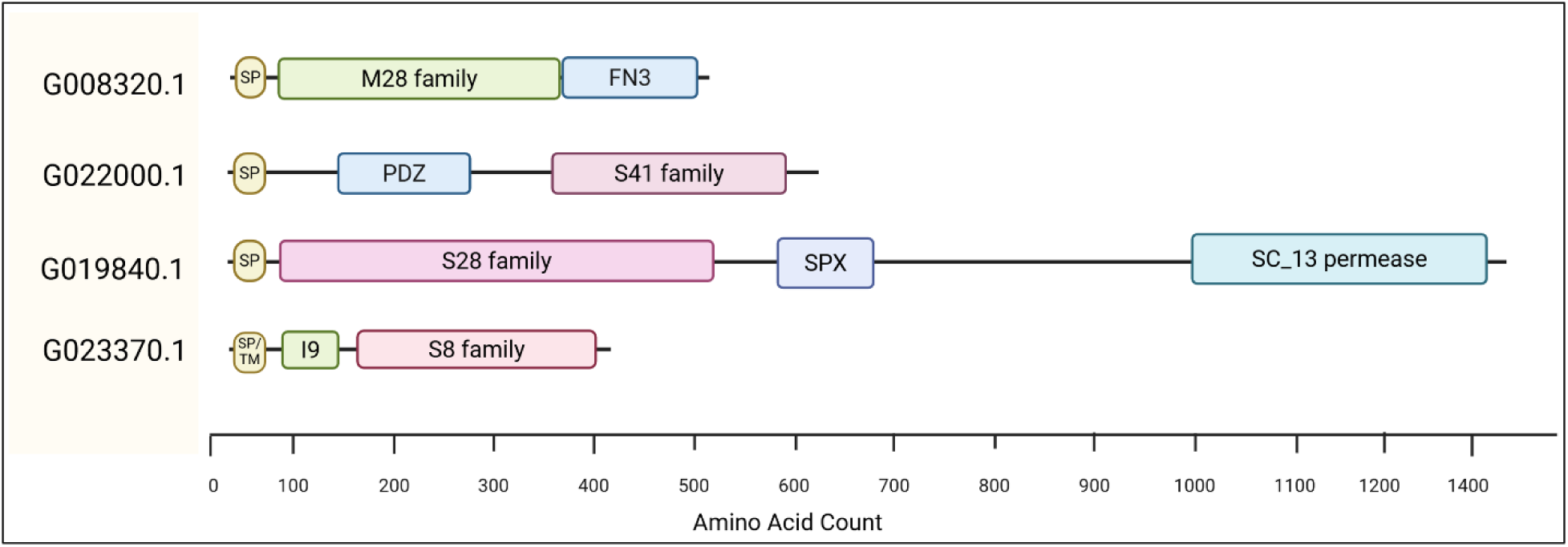
Domain architecture of multidomain proteases. Four multidomain proteases that were detected during growth on keratin-rich substrates and that are specifically discussed in the text are shown. The proteins are labeled with their accession number. The domains were assigned using Interpro Scan. The M28, S41, S28 and S8 domains belong to protease families, as classified in the MEROPS database. Abbreviations: I9, Peptidase Inhibitor I9; FN3, Fibronection III domain; PDZ, domain originally identified in PSD95, Dlg, and ZO-1 proteins; SPX, domain originally identified in SYG1, Pho81, and XPR1 proteins; SC_13 permease, solute carrier 13 permease; SP, signal peptide; SP/TM, a signal peptide and 1 transmembrane helix.

Clusters III and V include proteases belonging to the S8, S53, A1, M36, M35, M43 and M28 families, some of which were only detected during the growth on wool meal (Cluster III) and some only during growth on feather meal (Cluster V). Notably, the detection of these proteins, on substrates enriched in α-or β-keratin respectively, suggests that *O. corvina* tailors its proteolytic arsenal to accommodate distinctive biochemical properties of the substrates.

Interestingly, Cluster IV includes two proteases with transmembrane domains, an *endo*-protease from the S8 family (G023370.1) and an *exo*-protease from the S28 family (G019840.1). According to TMHMM 2.0 predictions, their catalytic domains are extracellularly oriented, suggesting a role in the breakdown of keratin-derived peptides before uptake by the fungal cell. Notably, the S28 family protease (G019840.1) also contains an SPX (SYG1/Pho81/XPR1) domain and a permease domain (Figure 4). SPX domains (IPR004331) have been implicated in G-protein signal transduction and phosphate sensing (40). This domain structure suggests that G019840.1 may have multiple roles, including peptide processing, transport, and signal transduction.

Altogether, the detection of many secreted proteases shows that *O. corvina* employs a comprehensive proteolytic system for keratin degradation (Figure 5). Cluster I proteases likely function as a general proteolytic system, hydrolyzing a broad range of proteins and peptides, while specialized proteases in other clusters target native keratin structures. The exclusive detection of Cluster III and V proteases in the wool and feather secretomes, respectively, shows that *O. corvina* tailors its enzymatic repertoire based on substrate composition and structure. By deploying a combination of both general and specific *endo*- and *exo*-proteases, possibly along with domain-specific adaptations that enhance substrate recognition and facilitate extracellular processing and peptide transport, the fungus ensures efficient keratin degradation, reinforcing its ability to thrive in keratin-rich environments.

**Figure 5:**
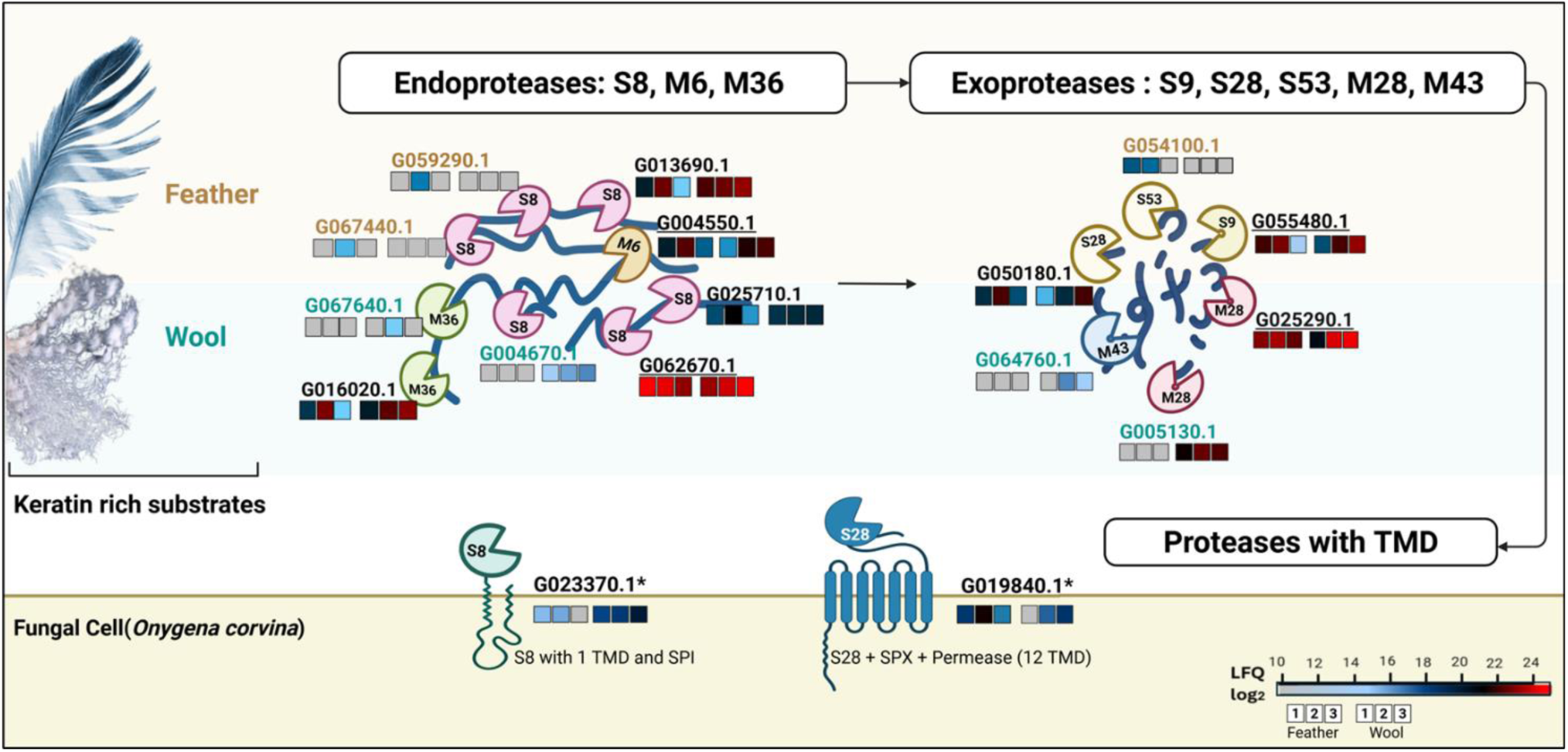
Proteolytic degradation machinery of *O. corvina*. The figure shows detected *endo*-proteases, *exo*-proteases and proteases with transmembrane domains (TMD involved in proteolytic degradation of keratin. Each detected protein is indicated with its accession number. The assigned peptidase family is indicated for each protease, based on the classification in the MEROPS database. Heat maps above or below the enzymes represent log_2_ LFQ intensities for proteins detected in the feather (left) and/or wool (right) secreteomes at days 1, 2, and 3. Table S2 lists the LFQ values for all detected putatively secreted proteases. The picture shows proteases detected across all substrates (Cluster I in Figure 3), those specific to keratin-rich substrates (Cluster II), and those primarily detected on feather meal (Cluster V; except G054750.1 and G047530.1) or wool meal (Cluster III), and those with predicted transmembrane domains. These latter two proteins belong to cluster IV (Fig. 3), which contains multiple low-abundance proteases that were detected in both the feather and wool secretomes, and that are not shown. Protein identifiers belonging to Clusters I and II are indicated in black, with those in Cluster I underlined. Identifiers corresponding to Clusters III and V are shown in blue and brown, respectively. Additionally, proteases with TMD have been highlighted with an asterisk (*).

### Expression of proteins involved in destabilizing keratin

Recent phylogenetic studies have positioned *O. corvina* within the dermatophytic *Arthrodermataceae* family (41, 42), suggesting that it may share keratin degradation strategies with dermatophytes. Given this classification, we speculated that the detection of proteins other than proteases could be associated with mechanisms akin to adherence and invasion of skin keratin by dermatophytes (17). These additional proteins may play a role in destabilizing the keratin structure, thereby enhancing proteolytic access. Based on the functional diversity of the detected proteins, this destabilization appears to occur through three complementary mechanisms that are discussed below: (1) mechanical disruption of keratin fibrils, (2) removal of post-translational modifications that reinforce keratin stability, and (3) removal of disulfide and other covalent bonds that provide structural rigidity (Figure 6).

**Figure 6:**
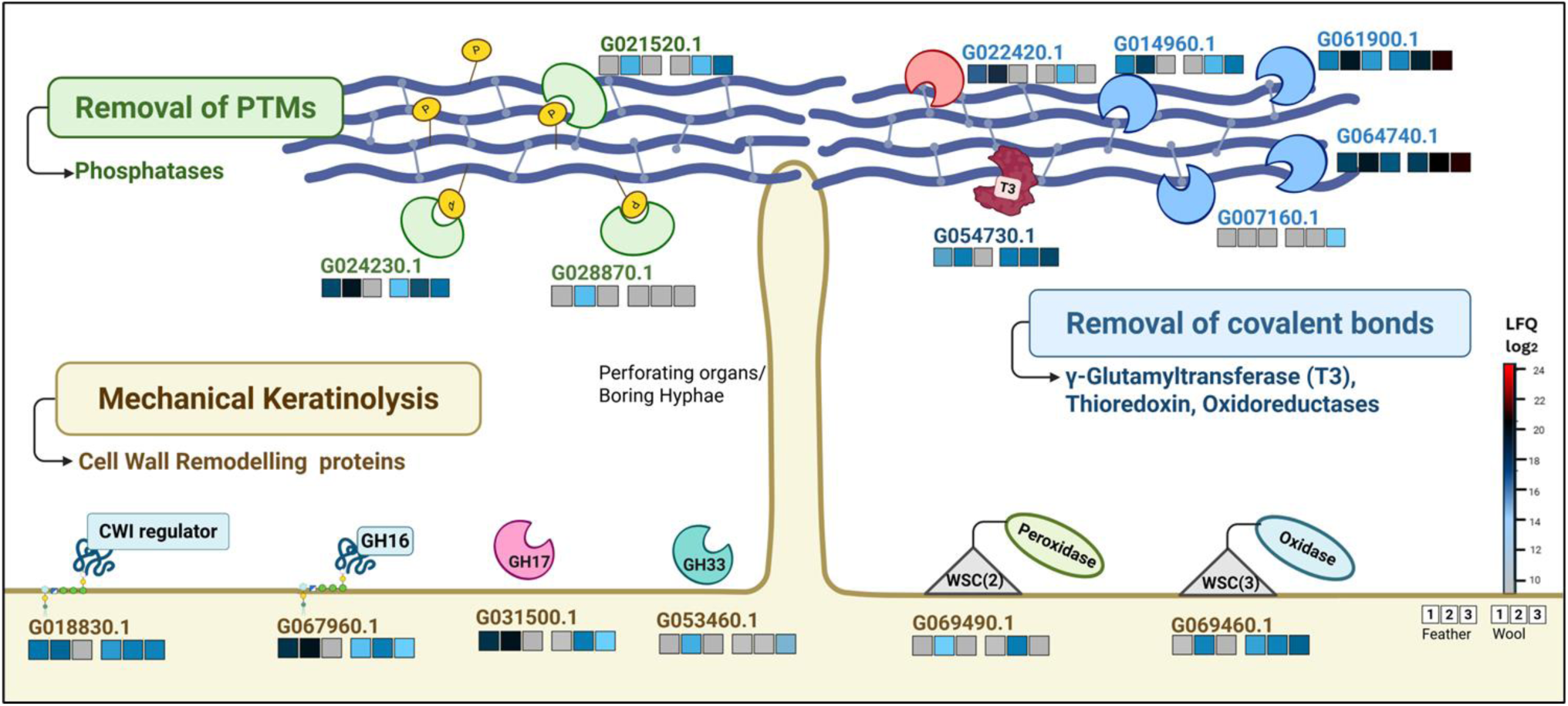
Keratin destabilization by *O. corvina*. The figure shows detected proteins involved in three complementary mechanisms of keratin destabilization: mechanical keratinolysis, removal of post-translational modifications, and removal of covalent bonds. Each protein is indicated with its accession number. Heat-maps above or below the enzymes represent log_2_ LFQ intensities for proteins detected in the feather (left) and/or wool (right) secretomes at days 1, 2, and 3. Table S2 lists the LFQ values for all the shown proteins, which are proteins involved in mechanical keratinolysis (G018830.1, G067960.1, G031500.1, G069490.1, G069460.1, G053460.1), phosphatases for removal of post-translational modifications (G021520.1, G024230.1, G028870.1), thioredoxin and oxidoreductases potentially involved in modifying disulfide bridges (G022420.1, G007160.1, G061900.1, G064740.1, G014960.1), and a T3 family gamma-glutamyl transpeptidase (G054730.1) involved in breaking iso-peptide bonds. The yellow blobs with a “P” represent phosphate groups; CWI, cell wall integrity; WSC, domains acting as mechanosensors that are typical for proteins involved in the CWI pathway.

### Mechanical Keratinolysis

Mechanical keratinolysis is a process in which certain fungi physically penetrate keratinized substrates using specialized structures like fronded mycelia or boring hyphae (43–45). These hyphae grow perpendicularly into the keratin matrix, facilitating colonization and subsequent enzymatic degradation. This phenomenon has been observed in fungi such as *Microsporum gypseum* (46) and *Trichophyton rubrum* (27), but the underlying molecular mechanisms remain largely unknown.

*O. corvina* mycelia formed dense aggregates with feather or wool meal after 8 hours of incubation in liquid minimal media. By 60 hours, fragmented substrate was released from these aggregates (Figure S6), suggesting keratin breakdown likely mediated by mycelial adhesion. Notably, about 20% of the 73 putatively secreted proteins detected during growth on feather and wool meal are involved in cell wall assembly and remodelling (Figure 2B). None of these were observed during growth on casein, indicating a keratin-specific role (Figure S5). The fungal cell wall, primarily composed of glucans and chitin, is dynamic and undergoes extensive reorganization in response to environmental stimuli (47). The detected cell wall-active enzymes include glucosidases, transglycosylases and sialidases, that are known to be involved in cell wall synthesis and remodeling and that may facilitate formation of mycelial structures that adhere to and mechanically disrupt keratin.

One such protein, G067960.1, is a GPI-lipid-anchored cell wall protein that is functionally annotated as a CRH1 transglycosylase of the Glycosyl hydrolase (GH16) family (as defined in the CAZy database (48)). In *Saccharomyces cerevisiae*, CRH1 crosslinks β-1,3-glucan and β-1,6-glucan to chitin and localizes at sites of polarized growth, such as budding tips, reinforcing the cell wall under stress conditions (49, 50). Given its functional annotation and sequence similarity (51% sequence identity), the *O. corvina* CRH1 ortholog may have a comparable role, potentially strengthening the fungal cell wall during keratin penetration and degradation. Another protein, G031500.1, belongs to the GH17 family and is involved in β-1,3-glucan modification. It is similar to EglC from *Aspergillus* spp., which is an *endo*-1,3-β-glucosidase that is highly conserved in fungi and has a known role in cell wall remodeling associated with hyphal extension (51–53). Recent studies have indicated that members of the GH16 and GH17 families may function collaboratively as glucan and chitin transferases for cell wall biogenesis (54), supporting the hypothesis that cell-wall remodelling is important during growth of *O. corvina* on keratin. Sialidases (G053460.1; GH33 family) have been implicated in fungal cell wall stability, particularly under mechanical stress conditions. In *Aspergillus fumigatus*, deletion of a sialidase encoding gene resulted in increased chitin deposition and unusual hyphal morphology, highlighting the role of this enzyme in maintaining cell wall integrity (55).

Additionally, the so-called Cell Wall Integrity (CWI) pathway is critical for responding to mechanical stress (56). Proteins involved in this pathway are often equipped with so-called WSC domains acting as mechanosensors (57). Two of the detected secreted proteins (G069460.1, G069490.1) were found to contain such WSC sensor domains. Another detected protein, G018830.1 is homologous to *ECM33*, a GPI anchored cell wall protein that is involved in the CWI pathway (58). The detection of these proteins suggests that *O. corvina* perceives keratin as mechanical stress, activating the CWI pathway in response.

The exclusive presence of proteins involved in cell wall synthesis and remodeling (GH16, GH17, and sialidases) along with cell wall integrity regulators (CWI pathway proteins) in the keratin secretomes, but not in the casein secretome, suggests involvement in mechanical keratinolysis.

### Removal of post translational modifications

Post-translational modifications (PTMs) such as phosphorylation, sumoylation, and glycosylation play a crucial role in maintaining the structure and stability of keratin, the conformation of which is regulated by its PTM status (19, 59). The removal of PTMs may contribute to destabilizing keratin and, indeed, multiple enzymes putatively involved in removing PTMs were detected in the secretome of *O. corvina* growing on feather or wool meal (Figure 6).

The phosphorylation of specific keratin-associated proteins has been implicated in the regulation of wool fiber crimping (60). These proteins form a matrix embedding keratin intermediate filaments that are cross-linked via disulphide bonds (61). We detected three phosphatases (G021520.1, G024230.1, G028870.1) annotated as alkaline phosphatase (IPR001952), histidine acid phosphatase (IPR016274) and calcineurin-like phosphoesterase (IPR014485), that could likely be involved in dephosphorylation of these keratin-associated proteins. Additionally, detected enzymes such as a β-hexoaminidase (G019890.1), a mannosidase (G049780.1), and two esterases (G036000.1, G042120.1) may be involved in modifying the glycosyl moieties found especially on the head domain of keratin polypeptide monomers. Such modifications will alter the nature of the filament head structures, ultimately leading to the disassembly of keratin filaments (13).

### Removal of covalent bonds

Lastly, another key mechanism contributing to keratin destabilization is the removal of covalent bonds such as disulfide bonds and keratin-specific isopeptide bonds (Figure 6). The secretomes of keratin-grown *O. corvina* contained several enzymes potentially involved in these processes, including a γ-glutamyl transferase (G054730.1), a thioredoxin (G022420.1) and four distinct FAD-containing oxidoreductases (G007160.1, G061900.1, G064740.1, G014960.1). None of these proteins were detected during growth on casein.

Based on previous studies on keratin degradation by *Bacillus sp.* CH1, the γ-glutamyl transferase is expected to cleave the isopeptide bonds between ε-amino groups of lysines and γ-glutamyl groups of glutamines, which commonly occur in keratinous proteins (62, 63). For disulfide bond reduction, the thioredoxin system has been widely studied. In this pathway, thioredoxins are maintained in their active redox state by FAD-containing thioredoxin reductases (64, 65). It is conceivable that the detected thioredoxin and FAD-containing oxidoreductases contribute to the reduction of disulfide bonds in keratin.

Taken together, the findings described so far suggest that keratin deconstruction in *O. corvina* is a multifaceted process integrating both mechanical and enzymatic processes (Figure 7). The fungus appears to employ a largely conserved strategy across α- and β-keratin-rich substrates, leveraging a combination of specific and generalized responses. On the one hand, targeted secretion of enzymes such as γ-glutamyl transferase will lead to destabilization of the keratin, while on the other hand, broader cellular stress responses, such as activation of the CWI pathway, will contribute to fungal robustness during the interaction with keratin and, possibly, mechanical disruption. This intricate balance between specific enzymatic action and global stress adaptation likely enables *O. corvina* to efficiently degrade keratinous substrates in diverse environmental contexts.

**Figure 7:**
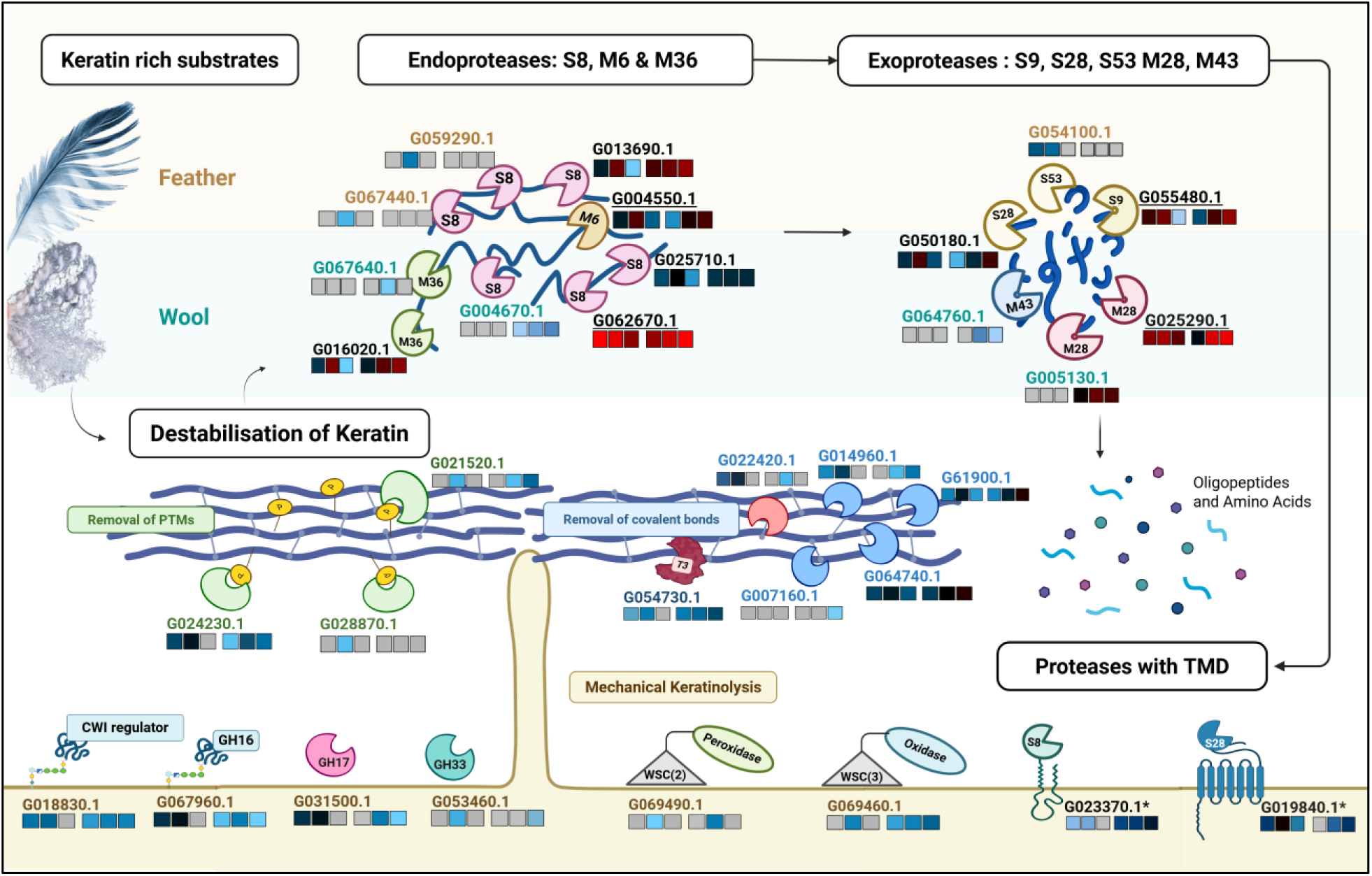
Overview of keratin degradation by *O. corvina*. Model for the degradation of keratin-rich substrates, such as feather and wool meal, by *O. corvina*. Destabilization of keratin includes mechanical keratinolysis (cell wall modifying enzymes), removal of PTMs (phosphatases) and removal of covalent bonds (glutamyl transpeptidase, thioredoxin and oxidoreductases). Proteolysis involves the action of *endo*-proteases, *exo*-proteases and proteases with transmembrane domains. Each protein is labelled with their accession number. See the legends of Fig. 5 C 6 for further explanations of the heat maps, colour coding and abbreviations.

### Substrate-specific keratinolytic response

As discussed above, *O. corvina* possesses a keratinolytic system capable of degrading both feather and wool substrates. The system contains many proteins that are used on both substrates, whereas each substrate specifically induces a small set of distinct secreted proteases. Feather meal stimulated production of proteases in Cluster V, whereas wool meal promoted production of proteases in Cluster III (Figure 3). This substrate-specific induction is similar to regulatory mechanisms observed in lignocellulose-degrading filamentous fungi, in which production of specific cellulases is driven by substrate-derived inducers (66, 67).

To further investigate whether *O. corvina* modulates its keratinolytic response in a substrate-dependent manner, we assessed its substrate degradation efficiency when cultivated on individual versus combined keratin-rich substrates. Given the variation in proteolytic machineries induced by these substrates and the fact that *O. corvina* inhabits decaying environments where it may encounter both α- and β-keratin forms, we hypothesized that the combination of substrates would elicit a broader and more synergistic enzymatic response. Comparative analyses revealed that when cultivated on a combined feather and wool meal substrate, *O. corvina* achieved 95% degradation after five days of incubation at 25 °C, a notable increase compared to approximately 70% degradation observed when grown on either feather or wool meal alone (Figure 8).

**Figure 8:**
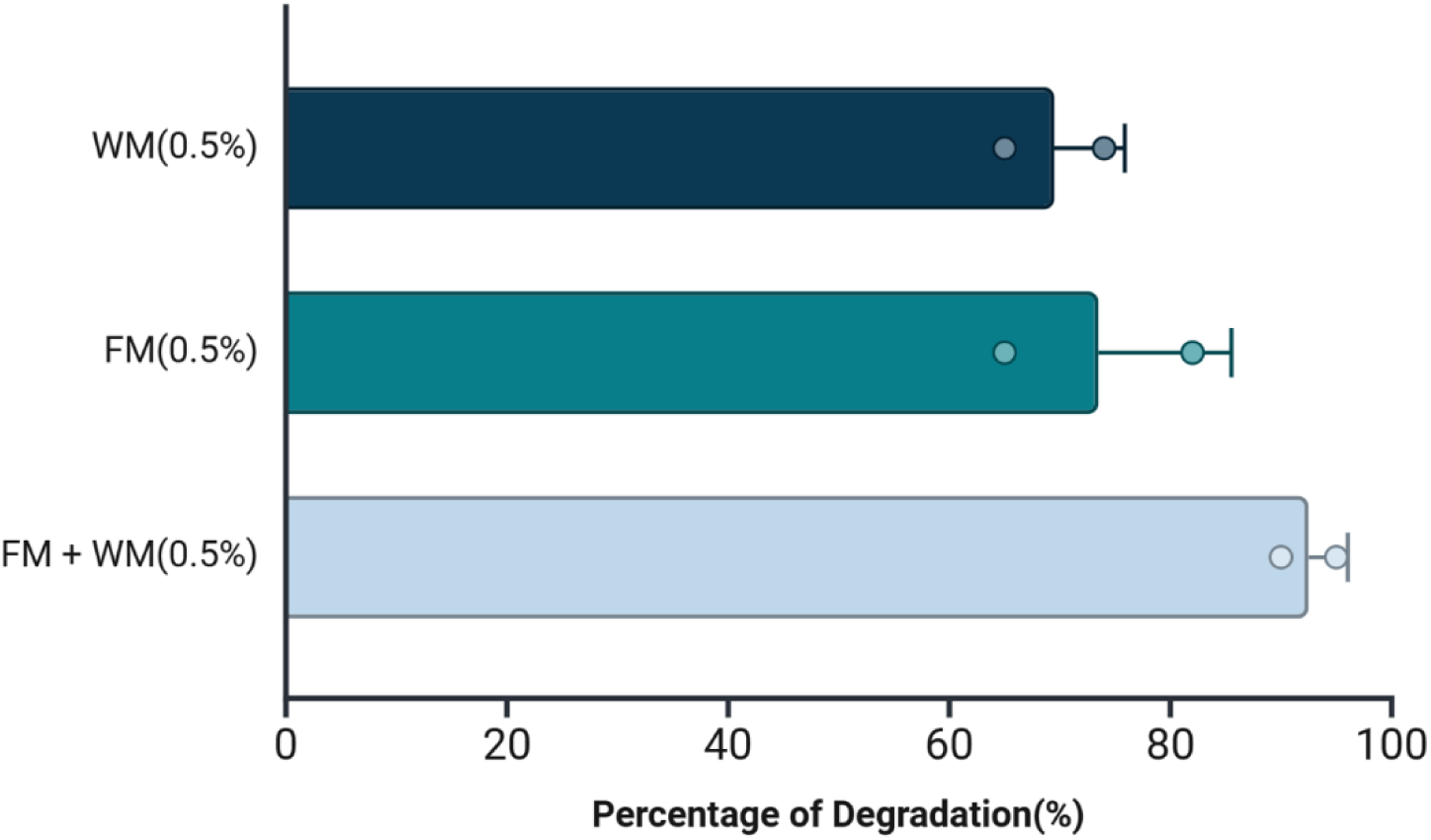
Degradation efficiency of feather meal, wool meal, and a combination thereof after 5 days of growth. Substrate degradation was measured by the weight loss method in cultures containing 0.5% (w/v) feather meal (FM), 0.5 % (w/v) wool meal (WM), or 0.25 % (w/v) of each of these two (FM+WM).

Proteolytic and keratinolytic enzyme assays showed that the supernatant from the combined feather and wool culture had the highest enzyme activity, compared to the supernatants of the cultures with individual substrates (Figure S7). This increase in activity correlates with the observed enhancement in degradation efficiency and suggests that the proteolytic machineries induced by different keratin-rich substrates act synergistically.

### Concluding remarks

In this work, we have used a proteogenomic approach to gain insights into the synergistic mechanisms employed by *O. corvina* for degrading keratin-rich materials. We focused on β- and α-keratin-rich substrates, specifically feather and wool meal, and utilized agar plates to ensure enrichment of substrate-bound and secreted proteins in the secretomes. This latter method simulates the natural solid-state degradation of keratin substrates by keratinolytic organisms, enabling the detection of relevant proteins involved in the process, while minimizing (commonly observed) contamination with non-secreted proteins. Indeed, the secretomes discussed above were strongly enriched in proteins that are predicted to be secreted.

Our data show that *O. corvina* employs a complex keratin degradation mechanism. Next to a extensive proteolytic machinery, the fungus likely destabilizes keratin through mechanical keratinolysis, removal of post-translational modifications, reduction of disulfide bonds, and cleavage of iso-peptide bonds, all of which enhance keratin accessibility for subsequent proteolysis. This process is facilitated by specific enzymatic activities (such as phosphatases and oxidoreductases) and a general stress response (including the CWI pathway). Proteolysis involves a combination of *endo*- and *exo*-proteases, several of which may have specific adaptations, in the form of additional domains, for substrate selectivity and peptide transport. As a consequence of these multiple mechanisms, keratin is converted into peptides and amino acids, making this recalcitrant material bioavailable for fungal uptake (Figure 7). Our data also shows that *O. corvina* to some extent adapts its enzyme repertoire to the nature of the keratinous substrate, suggesting that distinct keratin compositions provide complementary biochemical cues for the enzymatic response.

These novel insights into microbial degradation of keratin may guide the development of more efficient enzyme cocktails or microbial strains for keratin valorization. The complexity of the enzymatic machinery uncovered in this study indicates that development of efficient enzyme cocktails is challenging, perhaps pointing at microbial conversion as the preferred way to go. In this respect, *O. corvina* and its keratinolytic system provide a promising starting point. Further studies of individual members of the diverse enzymatic repertoire of *O. corvina*, including proteases, oxidoreductases, esterases, phosphatases, and sialidases, are of interest, since these enzymes hold biotechnological potential. Ultimately, the present findings emphasize the need for a holistic approach in designing biotechnological solutions for processing keratin-rich byproducts, advancing keratin valorization, and promoting sustainable byproduct management.

## Materials and Methods

### Strain and growth conditions

*O. corvina* (strain number: CBS 281.48; CBS-KNAW Collection of the Westerdijk Fungal Biodiversity Institute, Utrecht, Netherlands) was cultured on potato dextrose agar (PDA; Sigma-Aldrich, St. Louis, MO, USA) plates and subsequently cultivated on minimal medium plates supplemented with 1% (w/v) feather meal, wool meal, or casein. The composition of the minimal medium was as follows: 2 g/L magnesium sulfate heptahydrate (MgSO_4_ × 7H_2_O), 0.1 g/L potassium dihydrogen phosphate (KH_2_PO_4_), 0.01 g/L iron(II) sulfate heptahydrate (FeSO_4_ × 7H_2_O), 0.13 g/L calcium chloride dihydrate (CaCl_2_ × 2H_2_O), 10 g/L feather meal/wool meal/casein, and 15 g/L agar (VWR®, Radnor, PA, USA). Casein (C7078) was purchased from Sigma-Aldrich (St. Louis, MO, USA). Feather and wool meal were prepared separately using raw materials, provided by Norilia AS (Oslo, Norway). The materials were washed with warm running tap water until visually clean and then autoclaved at 130 °C, 3 atm for 40 minutes. This material was subsequently dried at 100 °C for 5 hours (68). The dried feathers were then ground using a bead beater with 50 mL stainless steel milling jars (Retsch, Haan, Germany) and 10 mm steel milling balls (VWR®, Radnor, USA), for 5 minutes at 20 rpm (69). The resulting fine powder was stored in an airtight container at room temperature until further use.

For the preparation of agar plates for secretome analysis, membrane plates were prepared as described by Bengtsson et al (70). Each plate consisted of two identical layers of the minimal medium supplemented with the respective substrate, which were separated by a sterile Supor 200 membrane (0.2 μm pore size, 47 mm diameter; Pall Life Sciences, Port Washington, USA). This membrane facilitates the separation of fungal cells, which stay in the upper agar layer, from the secreted enzymes that diffuse through the filter into the lower agar layer.

Plates containing the respective substrates and without membrane were inoculated by placing a 4 mm agar plug from potato dextrose agar plates, at the center of the plate. After incubation at 25 °C for two weeks, agar plugs from these plates were transferred to membrane-containing minimal agar plates containing the corresponding substrate. The plates were then incubated at 25 °C for 1, 2, and 3 days, after which samples for secretome analysis were collected.

### Sample Preparation for Proteomics

At each time point, agar discs from nine independent membrane plates (three biological replicates for each of the three substrates) were collected by punching out discs with an approximate volume of 100 µl from the bottom agar layer (70). Each disc was mixed with an equal volume of extraction buffer (10% SDS, 20 mM DTT, 100 mM Tris-HCl, pH 7.9) and incubated at 95 °C for 10 minutes to dissolve the agar and solubilize all proteins, also those bound to substrate. After vortexing and centrifugation (10 minutes at 5000×g), the supernatant was concentrated to 10–15 μL using vacuum centrifuge (SpeedVac, Thermo Fisher Scientific, Waltham, USA) and loaded on an SDS-PAGE gel, which was run at 220 mV for 2 minutes. Protein bands were stained with Coomassie brilliant blue, destained overnight, and excised. The gel pieces were washed, decolorized twice with 200 µl of 50% acetonitrile/25 mM ammonium bicarbonate, dehydrated by washing with 100µl of 100% acetonitrile, and air-dried before reduction (50µl of 10 mM DTT, 100 mM ammonium bicarbonate at 56 °C for 30 min) and alkylation (50µl of 55 mM iodoacetamide, 100 mM ammonium bicarbonate at room temperature in the dark for 30 min). Following removal of the excess alkylation solution, 200 μL of 100% acetonitrile was added to the gel pieces and incubated for 15 min at room temperature. After air drying, proteins in the gel pieces were digested overnight at 37 °C with 30 μL of 10 ng/ μL trypsin solution (Promega, Mannheim, Germany) (71). Digestion was stopped by adding 30 μL TFA to a final concentration of 0.5 % (v/v). After sonication for 15 minutes, the peptides were desalted using a STAGE-TIP protocol (72), dried under vacuum, and dissolved in 10 μL of 0.1% (v/v) formic acid.

### Proteomic analysis

The peptide samples were analyzed using a nano-UPLC (nanoElute 2, Bruker) coupled to a trapped ion mobility spectrometry/quadrupole time-of-flight mass spectrometer (timsTOF Pro, Bruker). Peptide separation was performed on an Aurora C18 reverse-phase analytical column (1.6 µm, 120 Å, 25 cm × 75 µm) with an integrated emitter (IonOpticks, Melbourne, Australia), maintained at 50°C using an integrated oven. The column was equilibrated at 800 bar before sample loading. Peptides were eluted at a flow rate of 300 nL/min using a solvent gradient from 5% to 25% solvent B over 40 min, followed by an increase to 37% over 5 min. The solvent composition was then raised to 95% solvent B over 5 min and maintained for an additional 10 min, resulting in a total run time of 60 min. Solvent A consisted of 0.1% (v/v) formic acid in Milli-Q water, while solvent B was 0.1% (v/v) formic acid in LC-MS grade acetonitrile. The timsTOF Pro was operated in positive ion data-dependent acquisition PASEF mode and controlled using Compass Hystar version 6.2.1.13 and timsControl version 5.0.4 (30538b33). The acquisition mass range was set to 100–1,700 m/z, with TIMS settings: 1/K0 Start 0.85 V⋅s/cm² and 1/K0 End 1.4 V⋅s/cm², Ramp time 100 ms, Ramp rate 9.42 Hz, Duty cycle 100%. The capillary voltage was set to 1,400 V, dry gas at 3.0 L/min, and dry temperature at 180 °C. The MS/MS settings included 10 PASEF ramps, a total cycle time of 0.53 s, a charge range of 0–5, a scheduling target intensity of 20,000, an intensity threshold of 2,500, active exclusion release after 0.4 min, and CID collision energy ranging from 27–45 eV.

Protein quantification was performed using the MSFragger v4.0 search engine (28, 73) within FragPipe v20.0, employing the LFQ-MBR workflow. A closed search (74) was conducted against the predicted *O.corvina* proteome (PRJNA1280007, 7232 proteins) supplemented with common contaminants and reversed decoy sequences for estimation of the false discovery rate (FDR). Label-free quantification with match-between-runs (MBR) was performed using IonQuant (75) v1.9.8. Peptide-spectrum matches (PSMs) were validated using Percolator (76), and protein inference was carried out with ProteinProphet(76). FDR filtration thresholds were set at 1% at both PSM and protein levels using Philosopher v5.0.0, employing a standard target-decoy database approach (78). Carbamidomethylation of cysteines was set as a fixed modification. Variable modifications included methionine oxidation, N-terminal acetylation, and pyro-glutamic acid formation at N-terminal glutamines. One missed cleavage was allowed. Normalization was applied in IonQuant to correct for technical variability and ensure comparability of protein abundance values across runs (74). The output was further processed in Perseus(79) v1.6.15.0. Proteins identified as potential contaminants, reverse hits, or proteins only identified by single peptide were excluded. Proteins were considered detected only if identified in at least two of three replicates. The LFQ intensities were log2 transformed prior to analysis. Hierarchical clustering and heatmap generation were done with Euclidian distance measure and complete linkage. Functional annotation of the detected proteins was then refined using InterPro Scan(80) version 5.73-104.0 and the MEROPS database(24). The subcellular localization of the detected proteins was predicted using a combination of SignalP 5.0 (81) and TMHMM (82) v2.0.

### Substrate Degradation and Enzyme Activity Assays

To assess substrate degradation dynamics, *O. corvina* was cultured in 300-ml Erlenmeyer flasks containing 100 ml minimal medium (composition described above) and 0.5 g of feather meal, wool meal, or a combination of feather and wool meal (each at 0.25 g). The flasks were inoculated with 1 g of wet mycelium. All flasks were incubated on a rotary shaker at 100 rpm and 25 °C for up to 5 days.

The experiment was performed in triplicate. Substrate degradation was monitored visually and recorded through images taken from the bottom of the conical flasks at 0, 8 and 60 hours. After 5 days of incubation, substrate degradation was evaluated using the weight loss method(82). The substrate-mycelial aggregate was collected onto a funnel lined with miracloth (Merck KGaA, Darmstadt, Germany), and the mycelial biomass was washed with 10% sodium hydroxide(84), until most of the adhering substrate was removed. The washed mycelial biomass was removed from the funnel with a sterile forcep, and the wet weight of the residual substrate was measured. The miracloth, containing the wet substrate, was then placed in a moisture analyzer (SARTORIUS MA160 Moisture Analyzer, Sartorius AG, Göttingen, Germany), where the moisture weight was determined. The final dry weight was calculated by subtracting the moisture weight from the wet weight of the residual substrate. The percentage of degradation was calculated using the formula:

*Percentage of Degradation (%) = [(0.5 – Dry weight of residual substrate) / 0.5] × 100*

Supernatants from feather meal, wool meal, combined substrate, and control cultures were diluted in 50 mM potassium phosphate buffer (pH 6.0) to normalize protein concentration across all samples. These protein-normalized supernatants were used to assess proteolytic and keratinolytic activities. Each reaction (200 µL total volume) contained 100 µL of the diluted supernatant and 100 µL of either a 2% (w/v) azocasein solution (for proteolytic activity; Megazyme, Bray, Ireland) or a suspension containing 2 mg of keratin azure (for keratinolytic activity; Sigma-Aldrich, St. Louis, USA), both prepared in the same buffer. Reactions were incubated at 37 °C for 120 minutes (proteolytic activity) or 24 hours (keratinolytic activity). For azocasein-containing samples, the reaction was terminated by adding 600 µL of 5% trichloroacetic acid (TCA; Sigma-Aldrich, St. Louis, MO, USA). All samples were centrifuged at 5000 × *g* for 10 minutes, and the supernatants were transferred to a 96-well plate. Absorbance was measured at 440 nm for proteolytic activity and at 595 nm for keratinolytic activity using a multimode microplate reader (Varioskan™ LUX, Thermo Fisher Scientific, Waltham, USA).

## Data availability

- The mass spectrometry data have been deposited to the ProteomeXchange Consortium via the PRIDE (85) partner repository with the dataset identifier PXD065899.
- The genome has been made available in the NCBI GenBank under the accession number PRJNA1280007. The gene and protein annotations, putative functional annotation, and CAZyme predictions are available on Figshare (https://doi.org/10.6084/m9.figshare.29328893.v1).

## Declarations

- Funding: This work was supported by the SFI Industrial Biotechnology program (project number 309558) funded by the Research Council of Norway.
- Authors’ contributions: SP: Performed experiments, analyzed data, and drafted the manuscript. CI: Prepared substrates and reviewed the manuscript. TRT: Guided experimental work and reviewed the manuscript. SLLR: Guided data analysis and curation; reviewed the manuscript. VS: Provided data insights; reviewed the manuscript. VE: Provided crucial insights; revised and reviewed the manuscript. All authors read and approved the final manuscript.
- Ethics approval and consent to participate: Not Applicable
- Competing interests: Not Applicable

## Supplemental Material

**Figure S1:**
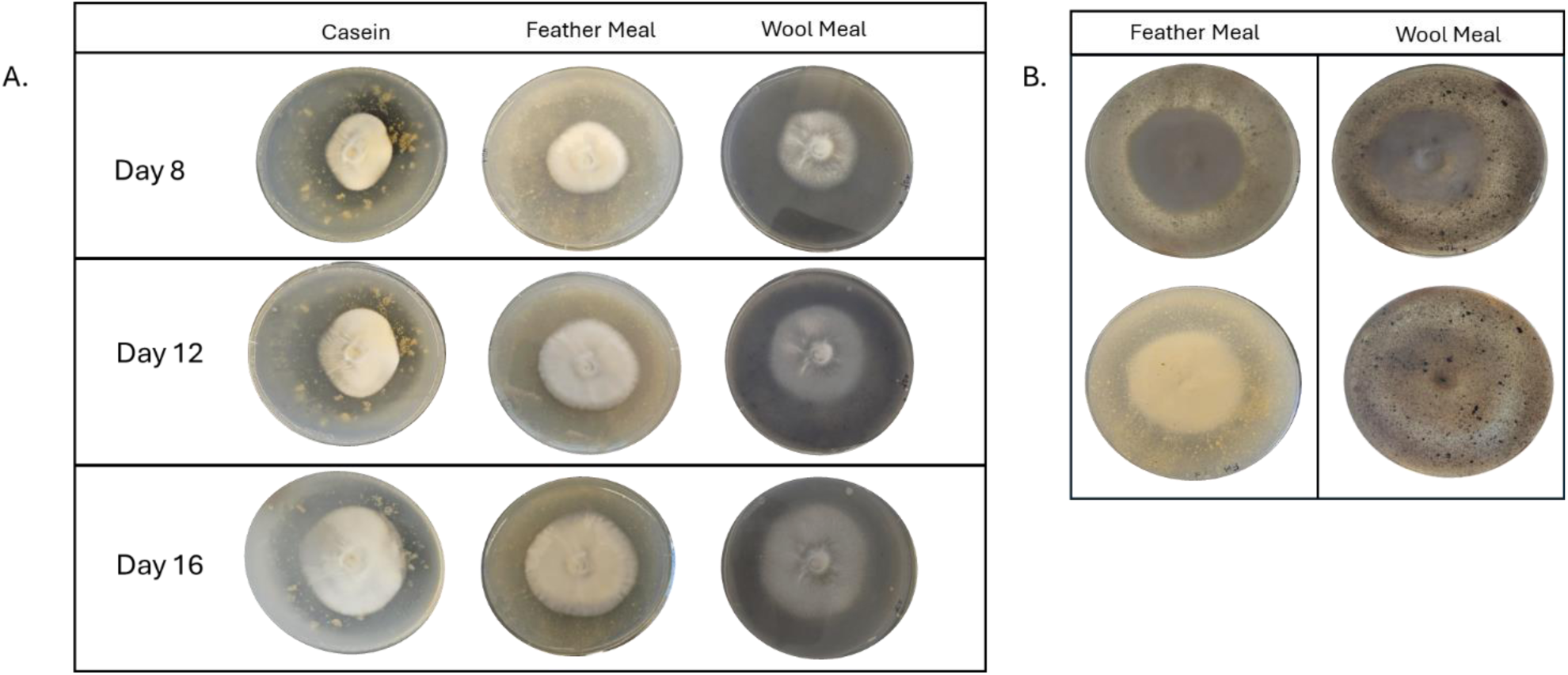
Growth and clearance zone of *O. corvina* growing on 1% casein, feather meal, or wool meal. (A) Radial growth of fungal colonies on minimal medium plates containing 1% casein, feather meal, or wool meal as the sole carbon and nitrogen source. Images were taken on days 8, 12, and 16 after incubation at 25 °C. A zone of clearance is visible around the colony on the casein plates. Due to the relatively low solubility of the keratin-rich substrates, clearance zones are less distinct for feather and wool meal. (B) To improve the visibility of clearance zones on feather and wool meal, the top (upper row) and bottom (lower row) views of plates at day 12 were photographed against bright light.

**Figure S2:**
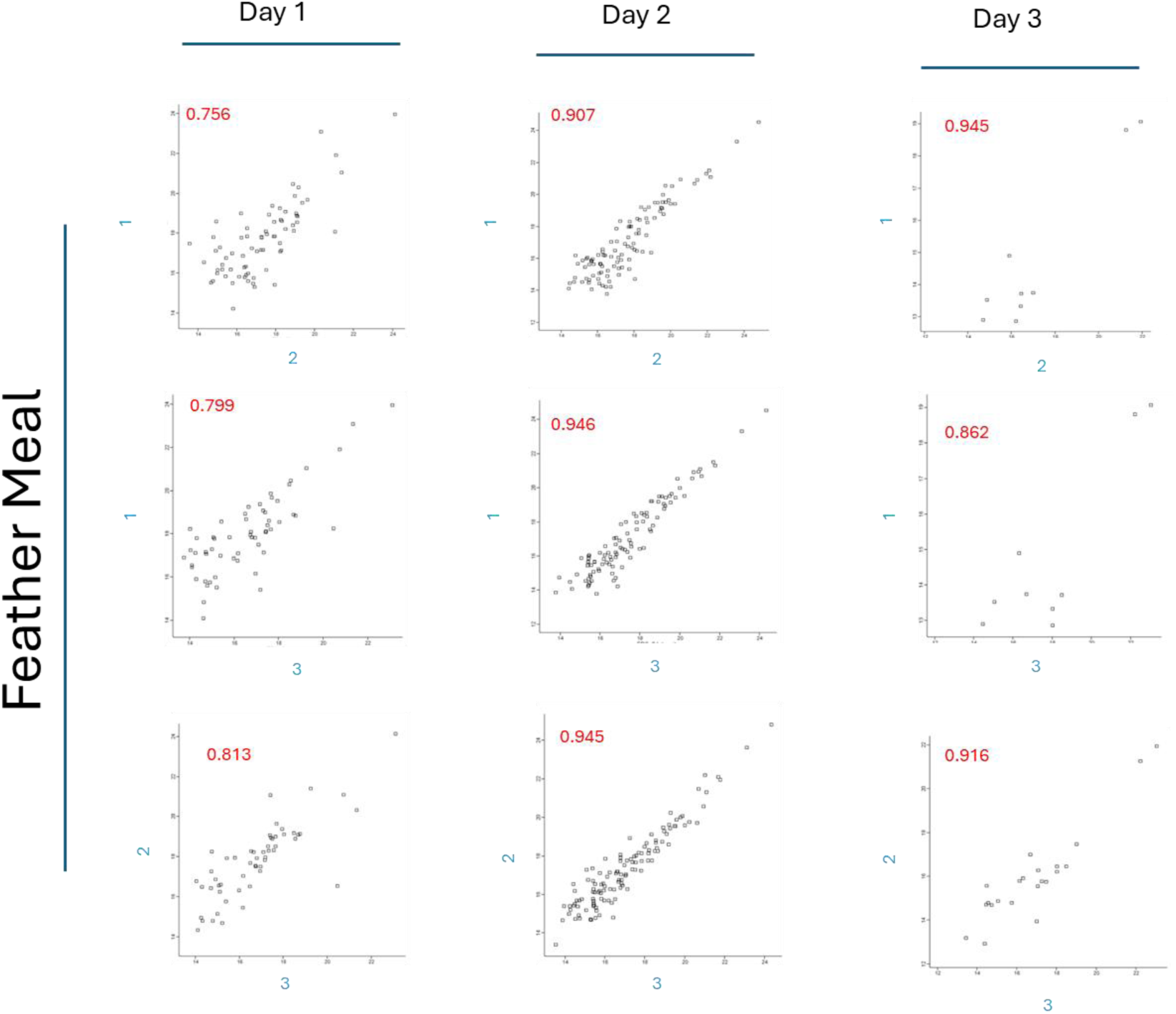
Multi-scatter plots showing pairwise comparisons of biological replicates for the feather secretome at Day 1, Day 2, and Day 3. Each scatter plot indicates the protein abundance values (log_2_ LFQ intensity) of two replicates at a given time point (Replicates 1–3), plotted on the x- and y-axes. Each dot represents a protein detected in both replicates. The Pearson correlation coefficient (r) between each replicate pair is displayed in red within each plot.

**Figure S3:**
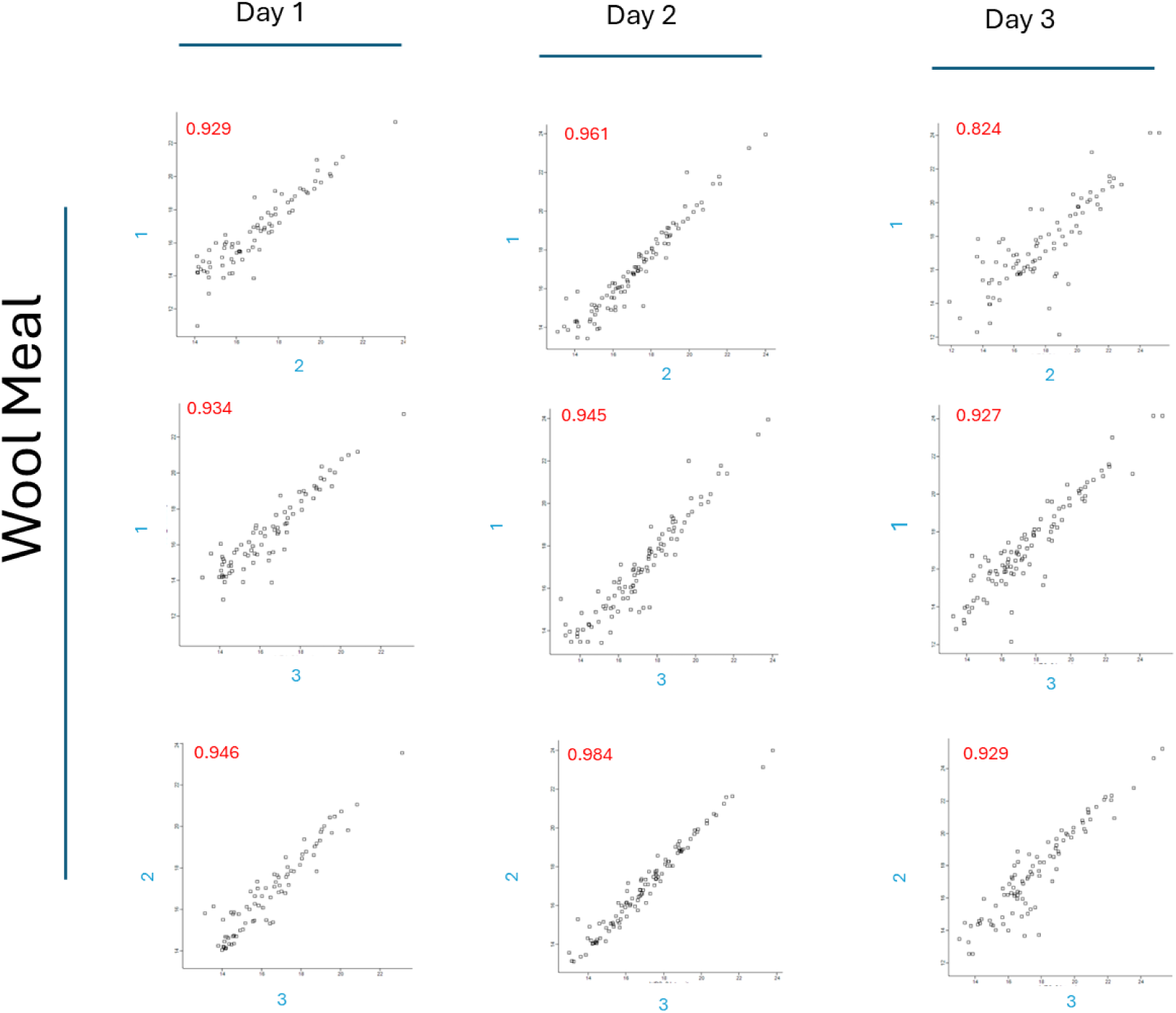
Multi-scatter plots showing pairwise comparisons of biological replicates for the wool secretome at Day 1, Day 2, and Day 3. Each scatter plot indicates the protein abundance values (log_2_ LFQ intensity) of two replicates at a given time point (Replicates 1–3), plotted on the x- and y-axes. Each dot represents a protein detected in both replicates. The Pearson correlation coefficient (r) between each replicate pair is displayed in red within each plot.

**Figure S4:**
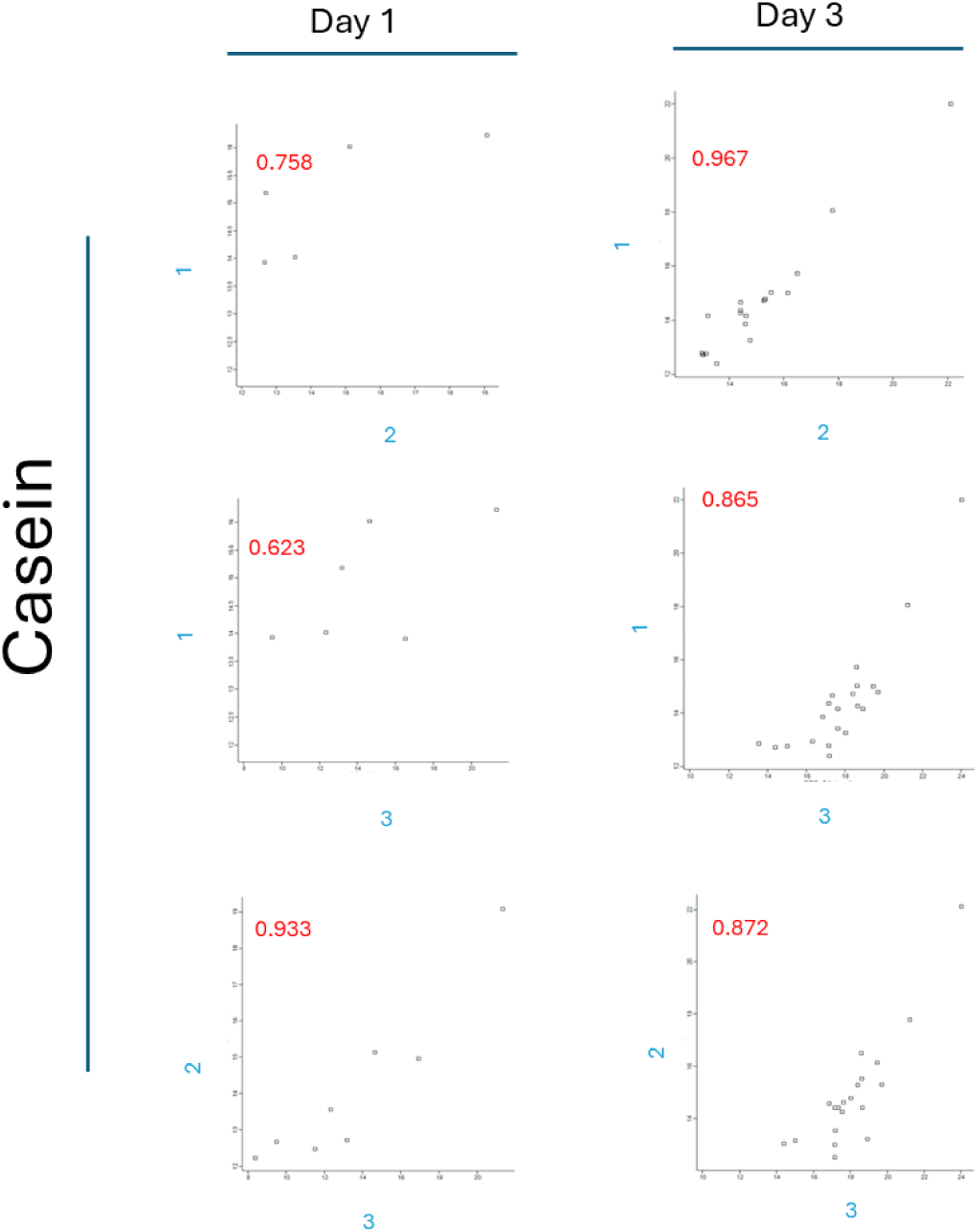
Multi-scatter plots showing pairwise comparisons of biological replicates for the casein secretome at Day 1, Day 2, and Day 3. Each scatter plot indicates the protein abundance values (log_2_ LFQ intensity) of two replicates at a given time point (Replicates 1–3), plotted on the x- and y-axes. Each dot represents a protein detected in both replicates. Since only one protein (a protease) was detected on day 2 with casein, this timepoint is not included in this analysis. The Pearson correlation coefficient (r) between each replicate pair is displayed in red within each plot.

**Figure S5:**
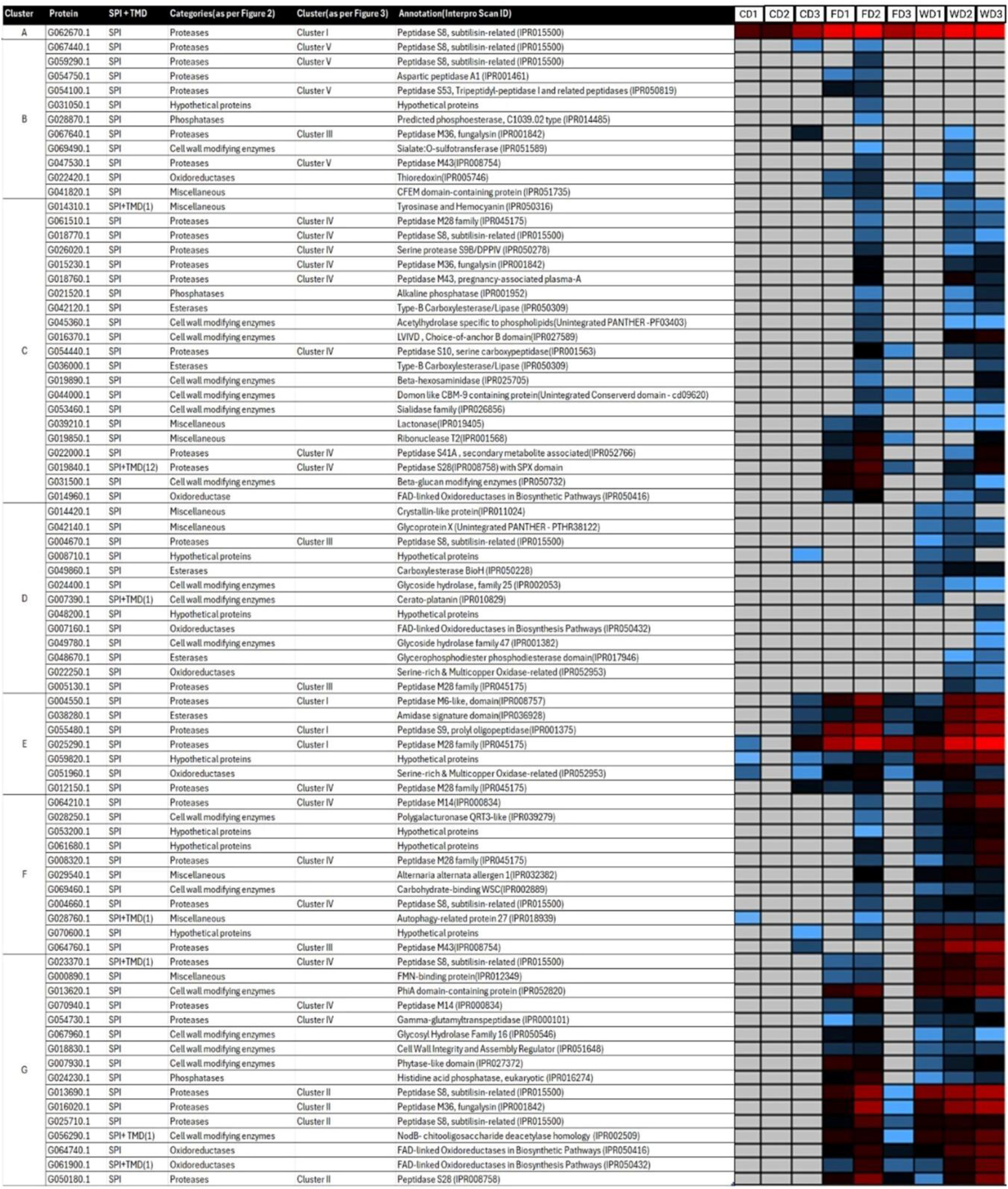
Functional annotation and heat map representation of 80 proteins (with predicted signal peptide – SPI) during growth on feather meal (FD1, FD2, FD3), wool meal (WD1, WD2, WD3), or casein (CD1, CD2, CD3) at day 1, 2, and 3. Each row represents a protein labeled with its accession number, and the color intensity reflects protein abundance (average of three replicates) at days 1, 2, or 3 on feather meal, wool meal, or casein. Note that seven of these proteins contain one or more transmembrane domains (indicated by “TMD”) and are thus not likely to be secreted. The heat map scale ranges from high abundance (red) to low abundance (light blue). Grey indicates proteins not detected. Proteins are divided into hierarchical clusters based on protein abundance (A, B, C, D, E, F, and G). The table also shows functional categories (as per Figure 2B), protease clusters (I-V; as per Figure 3), and annotations according to InterPro Scan (entry name and accession number, representing a protein superfamily, family, domain or repeat). In some cases, for entries lacking in InterPro, the PANTHER/CDD accession number is provided.

**Figure S6:**
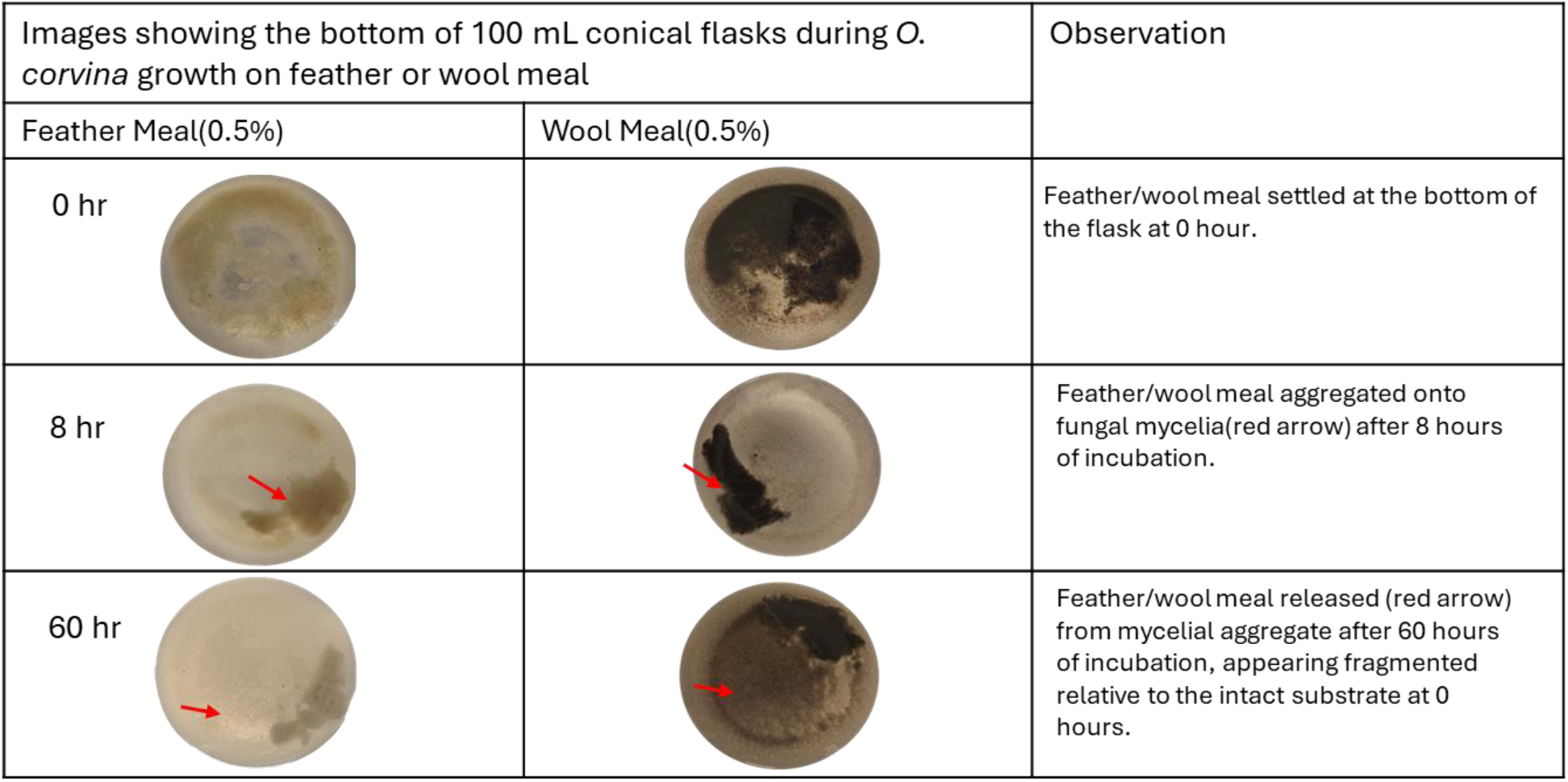
Live cell-substrate contact during growth of *O. corvina* on feather or wool meal. Images show the bottom of 100 mL conical flasks containing *O. corvina* cultures inoculated with 1 g of freshly cultivated mycelium in minimal media supplemented with 0.5% (w/v) feather meal or wool meal as the sole source of carbon and nitrogen. Cultures were incubated at 25 °C with shaking at 100 rpm, and images were captured at 0, 8, and 60 hours of incubation. The release of substrate from the mycelial aggregate at 60 hours is more clearly visible for wool meal than feather meal, because of better contrast against the background.

**Figure S7.**
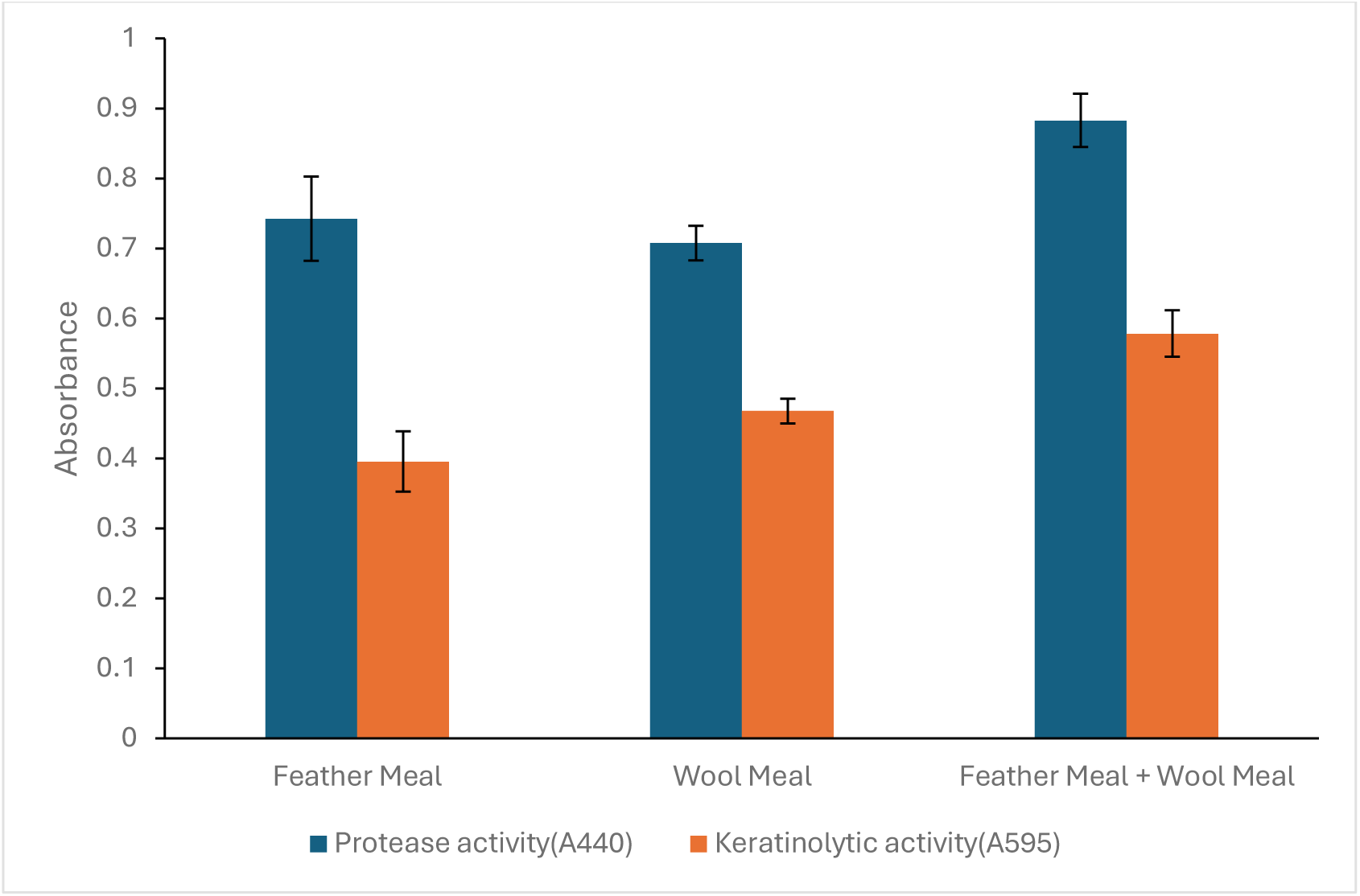
Proteolytic and keratinolytic activities in *O. corvina* cultures grown on feather meal, wool meal, and a combined substrate. Supernatants from *O. corvina* cultures grown for 5 days in feather meal (0.5%), wool meal (0.5%), or a combination of both (0.25% each) were tested for proteolytic and keratinolytic activities. Reactions were performed in 50 mM potassium phosphate buffer (pH 6.0) at 37 °C. Proteolytic activity was assessed using 1% azocasein, with absorbance measured at 440 nm after 120 minutes. Keratinolytic activity was measured using keratin azure (1% w/v), with absorbance recorded at 595 nm after 24 hours.

---

**Table S1 and S2 have been uploaded separately under ‘Other files’ due to their large size.**

**Table S1: Log₂ LFQ intensities and predicted signal peptide and transmembrane domains for all 154 detected proteins.**

Description of data: The Table contains the Log₂ LFQ intensities at day 1, 2, and 3 of cultivation for casein (CD1, CD2, CD3), feather meal (FD1, FD2, FD3), and wool meal (WD1, WD2, WD3). Each row contains the accession, the signal peptide (SPI) and transmembrane domain (TMD) prediction status and Log₂ LFQ intensities for each substrate at different timepoints (average of triplicates).

**Table S2: Log₂ LFQ intensities for 80 detected proteins with a predicted signal peptide.**

Description of data: The table contains the Log₂ LFQ intensities at day 1, 2, and 3 for casein (CD1, CD2, CD3), feather meal (FD1, FD2, FD3), and wool meal (WD1, WD2, WD3) secretomes. Each row contains the accession, the signal peptide (SPI) and transmembrane domain (TMD) prediction status, the assigned functional category (as per Figure 2B), a functional annotation based on InterPro Scan, and the Log₂ LFQ values. The InterPro Scan annotation includes the entry name and accession number, representing a protein superfamily, family, domain or repeat. In some cases, for entries lacking in InterPro, the PANTHER/CDD accession number is provided.

**Table S3:** Percentage of secreted proteins in the *O. corvina* genome and in the various substrate-specific secretomes. The table shows the total number of detected proteins, the number of putatively secreted proteins, and the corresponding percentage of secreted proteins for each of the casein, feather and wool secretomes. It also includes the average percentage of putatively secreted proteins across the three substrate-specific secretomes. For comparison, similar numbers are provided for the predicted proteome of *O.corvina*.

| Source | Total number of detected proteins | Total number of putatively secreted proteins | Percentage of secreted proteins (%) |  |  |
| --- | --- | --- | --- | --- | --- |
| Casein secretome (control) | 24 | 13 | 54 | 52.7 | Percentage of putatively secreted proteins across all three substrates(average) |
| Feather meal secretome | 118 | 59 | 50 |  |  |
| Wool meal secretome | 125 | 67 | 54 |  |  |
| Genome-predicted proteome | 7233 | 284 | 3.9 |  |  |

## References

1. Chukwunonso Ossai I, Shahul Hamid F, Hassan A. 2022. Valorisation of keratinous wastes: A sustainable approach towards a circular economy. Waste Management 151:81–104.

2. Suarato G, Contardi M, Perotto G, Heredia-Guerrero JA, Fiorentini F, Ceseracciu L, Pignatelli C, Debellis D, Bertorelli R, Athanassiou A. 2020. From fabric to tissue: Recovered wool keratin/polyvinylpyrrolidone biocomposite fibers as artificial scaffold platform. Materials Science and Engineering 116:111151.

3. Holkar CR, Jain SS, Jadhav AJ, Pinjari DV. 2018. Valorization of keratin based waste. Process Safety and Environmental Protection 115:85–98.

4. Vidmar B, Vodovnik M. 2018. Microbial Keratinases: Enzymes with Promising Biotechnological Applications. Food Technology and Biotechnology 56:312–328.

5. Tang Y, Guo L, Zhao M, Gui Y, Han J, Lu W, Dai Q, Jiang S, Lin M, Zhou Z, Wang J. 2021. A Novel Thermostable Keratinase from *Deinococcus geothermalis* with Potential Application in Feather Degradation. Applied Sciences 11:3136.

6. Sharma I, Pranaw K, Soni H, Rawat HK, Kango N. 2022. Parametrically optimized feather degradation by *Bacillus velezensis* NCIM 5802 and delineation of keratin hydrolysis by multi-scale analysis for poultry waste management. Sci Rep 12:17118.

7. Navone L, Speight R. 2018. Understanding the dynamics of keratin weakening and hydrolysis by proteases. PLoS ONE 13:e0202608.

8. Das S, Das A, Das N, Nath T, Langthasa M, Pandey P, Kumar V, Choure K, Kumar S, Pandey P. 2024. Harnessing the potential of microbial keratinases for bioconversion of keratin waste. Environ Sci Pollut Res 31:57478–57507.

9. Kang E, Jin H, La JW, Sung J, Park S, Kim W, Lee D. 2020. Identification of keratinases from *Fervidobacterium islandicum* AW-1 using dynamic gene expression profiling. Microbial Biotechnology 13:442–457.

10. Ward HM. 1997. VIII. *Onygena equina*, willd., a horn-destroying fungus. Philosophical Transactions of the Royal Society of London Series B, Containing Papers of a Biological Character 191:269–291.

11. Mercer DK, Stewart CS. 2019. Keratin hydrolysis by dermatophytes. Medical Mycology 57:13– 22.

12. Qiu J, Wilkens C, Barrett K, Meyer AS. 2020. Microbial enzymes catalyzing keratin degradation: Classification, structure, function. Biotechnology Advances 44:107607.

13. Zhou B, Guo Y, Xue Y, Ji X, Huang Y. 2023. Comprehensive insights into the mechanism of keratin degradation and exploitation of keratinase to enhance the bioaccessibility of soybean protein. Biotechnol Biofuels 16:177.

14. Jana A, Halder SK, Dasgupta D, Hazra S, Mondal P, Bhaskar T, Ghosh D. 2020. Keratinase Biosynthesis from Waste Poultry Feathers for Proteinaceous Stain Removal. ACS Sustainable Chem Eng 8:17651–17663.

15. Rahayu S, Syah D, Thenawidjaja Suhartono M. 2012. Degradation of keratin by keratinase and disulfide reductase from *Bacillus* sp. MTS of Indonesian origin. Biocatalysis and Agricultural Biotechnology 1:152–158.

16. Yamamura S, Morita Y, Hasan Q, Yokoyama K, Tamiya E. 2002. Keratin degradation: a cooperative action of two enzymes from *Stenotrophomonas* sp. Biochemical and Biophysical Research Communications 294:1138–1143.

17. Baldo A, Monod M, Mathy A, Cambier L, Bagut ET, Defaweux V, Symoens F, Antoine N, Mignon B. 2012. Mechanisms of skin adherence and invasion by dermatophytes. Mycoses 55:218–223.

18. Eymann C, Wachlin G, Albrecht D, Tiede S, Krummrei U, Jünger M, Hecker M, Daeschlein G. 2018. Exoproteome Analysis of Human Pathogenic Dermatophyte Species and Identification of Immunoreactive Proteins. Proteomics Clinical Apps 12:1800007.

19. Lange L, Huang Y, Busk PK. 2016. Microbial decomposition of keratin in nature—a new hypothesis of industrial relevance. Appl Microbiol Biotechnol 100:2083–2096.

20. Huang Y, Busk PK, Herbst F-A, Lange L. 2015. Genome and secretome analyses provide insights into keratin decomposition by novel proteases from the non-pathogenic fungus *Onygena corvina*. Applied Microbiology and Biotechnology 99:9635–9649.

21. Huang Y, Busk PK, Lange L. 2015. Production and Characterization of Keratinolytic Proteases Produced by *Onygena corvina*. Fungal Genom Biol 05.

22. Nesvizhskii AI. 2014. Proteogenomics: concepts, applications and computational strategies. Nat Methods 11:1114–1125.

23. Pavale S, Eijsink VGH, La Rosa SL. 2025. Improved Genome Assembly of *Onygena corvina*, a Keratin-Degrading Fungus. bioRxiv 2025.07.02.662733.

24. Huang Y, Łężyk M, Herbst F-A, Busk PK, Lange L. 2020. Novel keratinolytic enzymes, discovered from a talented and efficient bacterial keratin degrader. Sci Rep 10:10033.

25. Rawlings ND, Barrett AJ, Bateman A. 2010. MEROPS: the peptidase database. Nucleic Acids Research 38:D227–D233.

26. Malviya HK, Rajak RC, Hasija SK. 1992. Synthesis and regulation of extracellular keratinase in three fungi isolated from the grounds of a gelatin factory, Jabalpur, India. Mycopathologia 120:1–4.

27. Marchisio V. 2000. Keratinophilic fungi: Their role in nature and degradation of keratinic substrates. Biology of Dermatophytes and Other Keratinophilic Fungi 17.

28. Teo GC, Polasky DA, Yu F, Nesvizhskii AI. 2021. Fast Deisotoping Algorithm and Its Implementation in the MSFragger Search Engine. J Proteome Res 20:498–505.

29. Rawlings ND. 2016. Peptidase specificity from the substrate cleavage collection in the MEROPS database and a tool to measure cleavage site conservation. Biochimie 122:5–30.

30. Lowther WT, Matthews BW. 2002. Metalloaminopeptidases: Common Functional Themes in Disparate Structural Surroundings. Chem Rev 102:4581–4608.

31. Skowron PM, Krefft D, Brodzik R, Kasperkiewicz P, Drag M, Koller K-P. 2020. An alternative for proteinase K-heat-sensitive protease from fungus *Onygena corvina* for biotechnology: cloning, engineering, expression, characterization and special application for protein sequencing. Microbial Cell Factories 19:135.

32. Qiu J, Barrett K, Wilkens C, Meyer AS. 2022. Bioinformatics based discovery of new keratinases in protease family M36. New Biotechnology 68:19–27.

33. Soisson SM, Patel SB, Abeywickrema PD, Bryne NJ, Diehl RE, Hall DL, Ford RE, Reid JC, Rickert KW, Shipman JM, Sharma S, Lumb KJ. 2010. Structural definition and substrate specificity of the S28 protease family: the crystal structure of human prolylcarboxypeptidase. BMC Struct Biol 10:16.

34. Koide A, Bailey CW, Huang X, Koide S. 1998. The fibronectin type III domain as a scaffold for novel binding proteins. Journal of Molecular Biology 284:1141–1151.

35. Van Leeuwen HC, Roelofs D, Corver J, Hensbergen P. 2021. Phylogenetic analysis of the bacterial Pro-Pro-endopeptidase domain reveals a diverse family including secreted and membrane anchored proteins. Current Research in Microbial Sciences 2:100024.

36. Muley VY, Akhter Y, Galande S. 2019. PDZ Domains Across the Microbial World: Molecular Link to the Proteases, Stress Response, and Protein Synthesis. Genome Biology and Evolution 11:644–659.

37. Fanning AS, Anderson JM. 1996. Protein–protein interactions: PDZ domain networks. Current Biology 6:1385–1388.

38. Ghosh A, Chakrabarti K, Chattopadhyay D. 2009. Cloning of feather-degrading minor extracellular protease from *Bacillus cereus* DCUW: dissection of the structural domains. Microbiology 155:2049–2057.

39. Zaugg C, Jousson O, Léchenne B, Staib P, Monod M. 2008. Trichophyton rubrum secreted and membrane-associated carboxypeptidases. International Journal of Medical Microbiology 298:669–682.

40. Duan K, Yi K, Dang L, Huang H, Wu W, Wu P. 2008. Characterization of a sub-family of Arabidopsis genes with the SPX domain reveals their diverse functions in plant tolerance to phosphorus starvation. Plant J 54:965–975.

41. Garcia Garces H, Hamae Yamauchi D, Theodoro RC, Bagagli E. 2020. PRP8 Intein in Onygenales: Distribution and Phylogenetic Aspects. Mycopathologia 185:37–49.

42. Kandemir H, Dukik K, de Melo Teixeira M, Stielow JB, Delma FZ, Al-Hatmi AMS, Ahmed SA, Ilkit M, de Hoog GS. 2022. Phylogenetic and ecological reevaluation of the order Onygenales. Fungal Diversity 115:1–72.

43. Peng Z, Zhang J, Du G, Chen J. 2019. Keratin Waste Recycling Based on Microbial Degradation: Mechanisms and Prospects. ACS Sustainable Chem Eng 7:9727–9736.

44. Błyskal B. 2009. Fungi utilizing keratinous substrates. International Biodeterioration C Biodegradation 63:631–653.

45. English MP. 1965. The saprophytic growth of non-keratinophilic fungi on keratinized substrata, and a comparison with keratinophilic fungi. Transactions of the British Mycological Society 48:219-IN8.

46. Kunert J, Krajcí D. 1981. An Electron Microscopy Study of Keratin Degradation by the Fungus Microsporum gypseum in vitro. Mycoses 24:485–496.

47. Ene IV, Walker LA, Schiavone M, Lee KK, Martin-Yken H, Dague E, Gow NAR, Munro CA, Brown AJP. 2015. Cell Wall Remodeling Enzymes Modulate Fungal Cell Wall Elasticity and Osmotic Stress Resistance. mBio 6:e00986–15.

48. Drula E, Garron M-L, Dogan S, Lombard V, Henrissat B, Terrapon N. 2022. The carbohydrate-active enzyme database: functions and literature. Nucleic Acids Research 50:D571–D577.

49. Cabib E, Farkas V, Kosík O, Blanco N, Arroyo J, McPhie P. 2008. Assembly of the Yeast Cell Wall. Journal of Biological Chemistry 283:29859–29872.

50. Cabib E, Blanco N, Grau C, Rodríguez-Peña JM, Arroyo J. 2007. Crh1p and Crh2p are required for the cross-linking of chitin to β(1-6)glucan in the *Saccharomyces cerevisiae* cell wall. Molecular Microbiology 63:921–935.

51. Ramirez-Garcia A, Pellon A, Buldain I, Antoran A, Arbizu-Delgado A, Guruceaga X, Rementeria A, Hernando FL. 2018. Proteomics as a Tool to Identify New Targets Against Aspergillus and Scedosporium in the Context of Cystic Fibrosis. Mycopathologia 183:273–289.

52. Liu J, Hu X. 2023. Fungal extracellular vesicle-mediated regulation: from virulence factor to clinical application. Front Microbiol 14:1205477.

53. Souza JAM, Baltazar LDM, Carregal VM, Gouveia-Eufrasio L, De Oliveira AG, Dias WG, Campos Rocha M, Rocha De Miranda K, Malavazi I, Santos DDA, Frézard FJG, De Souza DDG, Teixeira MM, Soriani FM. 2019. Characterization of *Aspergillus fumigatus* Extracellular Vesicles and Their Effects on Macrophages and Neutrophils Functions. Front Microbiol 10:2008.

54. Patel P, Free SJ. 2022. Characterization of Neurospora crassa GH16, GH17, and GH72 gene families of cell wall crosslinking enzymes. The Cell Surface 8:100073.

55. Nesbitt JR, Steves EY, Schonhofer CR, Cait A, Manku SS, Yeung JHF, Bennet AJ, McNagny KM, Choy JC, Hughes MR, Moore MM. 2018. The *Aspergillus fumigatus* Sialidase (Kdnase) Contributes to Cell Wall Integrity and Virulence in Amphotericin B-Treated Mice. Front Microbiol 8:2706.

56. Fuchs BB, Mylonakis E. 2009. Our Paths Might Cross: the Role of the Fungal Cell Wall Integrity Pathway in Stress Response and Cross Talk with Other Stress Response Pathways. Eukaryot Cell 8:1616–1625.

57. Oide S, Tanaka Y, Watanabe A, Inui M. 2019. Carbohydrate-binding property of a cell wall integrity and stress response component (WSC) domain of an alcohol oxidase from the rice blast pathogen *Pyricularia oryzae*. Enzyme and Microbial Technology 125:13–20.

58. Pardo M. 2004. PST1 and ECM33 encode two yeast cell surface GPI proteins important for cell wall integrity. Microbiology 150:4157–4170.

59. Bragulla HH, Homberger DG. 2009. Structure and functions of keratin proteins in simple, stratified, keratinized and cornified epithelia. Journal of Anatomy 214:516–559.

60. He D, Chen L, Luo F, Zhou H, Wang J, Zhang Q, Lu T, Wu S, Hickford JGH, Tao J. 2021. Differentially phosphorylated proteins in the crimped and straight wool of Chinese Tan sheep. Journal of Proteomics 235:104115.

61. Shi A, Ma S, Yang Z, Ding W, Tian J, Chen X, Tao J. 2024. Proteomic Analysis of Crimped and Straight Wool in Chinese Tan Sheep. Animals 14:2858.

62. Callegaro K, Brandelli A, Daroit DJ. 2019. Beyond plucking: Feathers bioprocessing into valuable protein hydrolysates. Waste Management 95:399–415.

63. Lai Y, Wu X, Zheng X, Li W, Wang L. 2023. Insights into the keratin efficient degradation mechanism mediated by Bacillus sp. CN2 based on integrating functional degradomics. Biotechnology for Biofuels and Bioproducts 16:59.

64. Buey RM, Fernández-Justel D, González-Holgado G, Martínez-Júlvez M, González-López A, Velázquez-Campoy A, Medina M, Buchanan BB, Balsera M. 2021. Unexpected diversity of ferredoxin-dependent thioredoxin reductases in cyanobacteria. Plant Physiology 186:285– 296.

65. Arnér ESJ, Holmgren A. 2000. Physiological functions of thioredoxin and thioredoxin reductase. European Journal of Biochemistry 267:6102–6109.

66. Novy V, Nielsen F, Seiboth B, Nidetzky B. 2019. The influence of feedstock characteristics on enzyme production in *Trichoderma reesei:* a review on productivity, gene regulation and secretion profiles. Biotechnol Biofuels 12:238.

67. Scott CJR, McGregor NGS, Leadbeater DR, Oates NC, Hoßbach J, Abood A, Setchfield A, Dowle A, Overkleeft HS, Davies GJ, Bruce NC. 2024. *Parascedosporium putredinis* NO1 tailors its secretome for different lignocellulosic substrates. Microbiol Spectr 12:e03943–23.

68. Strzetelski JA, Kowalczyk J, Niwińska B, Bilik K, Maciaszek K. 1999. Nutritive value of feather keratin meals for ruminants. J Anim Feed Sci 8:387–393.

69. Isembart C, Zimmermann B, Matić J, Bolaño Losada C, Afseth NK, Kohler A, Horn Svein J, Eijsink V, Chylenski P, Shapaval V. 2025. Comparative analysis of pre-treatment strategies and bacterial strain efficiency for improvement of feather hydrolysis. Microbial Cell Factories 24:118.

70. Bengtsson O, Arntzen MØ, Mathiesen G, Skaugen M, Eijsink VGH. 2016. A novel proteomics sample preparation method for secretome analysis of *Hypocrea jecorina* growing on insoluble substrates. Journal of Proteomics 131:104–112.

71. Shevchenko A, Tomas H, Havli J, Olsen JV, Mann M. 2006. In-gel digestion for mass spectrometric characterization of proteins and proteomes. Nat Protoc 1:2856–2860.

72. Rappsilber J, Mann M, Ishihama Y. 2007. Protocol for micro-purification, enrichment, pre-fractionation and storage of peptides for proteomics using StageTips. Nat Protoc 2:1896– 1906.

73. Kong AT, Leprevost FV, Avtonomov DM, Mellacheruvu D, Nesvizhskii AI. 2017. MSFragger: ultrafast and comprehensive peptide identification in mass spectrometry–based proteomics. Nat Methods 14:513–520.

74. Yu F, Haynes SE, Teo GC, Avtonomov DM, Polasky DA, Nesvizhskii AI. 2020. Fast Quantitative Analysis of timsTOF PASEF Data with MSFragger and IonQuant. Mol Cell Proteomics 19:1575– 1585.

75. Yu F, Haynes SE, Nesvizhskii AI. 2021. IonQuant Enables Accurate and Sensitive Label-Free Quantification With FDR-Controlled Match-Between-Runs. Molecular C Cellular Proteomics 20:100077.

76. Käll L, Canterbury JD, Weston J, Noble WS, MacCoss MJ. 2007. Semi-supervised learning for peptide identification from shotgun proteomics datasets. Nat Methods 4:923–925.

77. Nesvizhskii AI, Keller A, Kolker E, Aebersold R. 2003. A Statistical Model for Identifying Proteins by Tandem Mass Spectrometry. Anal Chem 75:4646–4658.

78. da Veiga Leprevost F, Haynes SE, Avtonomov DM, Chang H-Y, Shanmugam AK, Mellacheruvu D, Kong AT, Nesvizhskii AI. 2020. Philosopher: a versatile toolkit for shotgun proteomics data analysis. Nat Methods 17:869–870.

79. Tyanova S, Temu T, Sinitcyn P, Carlson A, Hein MY, Geiger T, Mann M, Cox J. 2016. The Perseus computational platform for comprehensive analysis of (prote)omics data. Nat Methods 13:731–740.

80. Jones P, Binns D, Chang H-Y, Fraser M, Li W, McAnulla C, McWilliam H, Maslen J, Mitchell A, Nuka G, Pesseat S, Quinn AF, Sangrador-Vegas A, Scheremetjew M, Yong S-Y, Lopez R, Hunter S. 2014. InterProScan 5: genome-scale protein function classification. Bioinformatics 30:1236–1240.

81. Almagro Armenteros JJ, Tsirigos KD, Sønderby CK, Petersen TN, Winther O, Brunak S, von Heijne G, Nielsen H. 2019. SignalP 5.0 improves signal peptide predictions using deep neural networks. Nat Biotechnol 37:420–423.

82. Krogh A, Larsson B, von Heijne G, Sonnhammer ELL. 2001. Predicting transmembrane protein topology with a hidden markov model: application to complete genomes1. Journal of Molecular Biology 305:567–580.

83. Li Z-W, Liang S, Ke Y, Deng J-J, Zhang M-S, Lu D-L, Li J-Z, Luo X-C. 2020. The feather degradation mechanisms of a new Streptomyces sp. isolate SCUT-3. Commun Biol 3:1–13.

84. Korniłłowicz-Kowalska T. 1997. Studies on the decomposition of keratin wastes by saprotrophic microfungi. I. Criteria for evaluating keratinolytic activity. 1. Acta Mycologica 32:51–79.

85. Perez-Riverol Y, Bandla C, Kundu DJ, Kamatchinathan S, Bai J, Hewapathirana S, John NS, Prakash A, Walzer M, Wang S, Vizcaíno JA. 2025. The PRIDE database at 20 years: 2025 update. Nucleic Acids Research 53:D543–D553.

